# Complement Dysregulation During the Early Phases of Synucleinopathy

**DOI:** 10.64898/2026.04.27.720696

**Authors:** Hina Khan, Mary Gifford, Arash Kordbacheh, Asher Bury, Spencer Panoushek, Allyson Cole-Strauss, Christopher J. Kemp, Kelvin C. Luk, Kathy Steece-Collier, Nathan C. Kuhn, Nicholas M. Kanaan, Caryl E. Sortwell, Joseph R. Patterson, Matthew J. Benskey

## Abstract

Parkinson’s disease (PD) is characterized by progressive degeneration of nigrostriatal dopamine neurons and synucleinopathy, which is the accumulation of aggregated α-synuclein (α-syn). Increasing evidence implicates α-syn-associated neuroinflammation as a contributor to PD pathogenesis; however, immune mechanisms linking synucleinopathy to neurodegeneration remain incompletely defined. Activation of the complement cascade occurs in PD and other neurodegenerative disorders, but most studies report complement activation after overt neurodegeneration, making it difficult to conclude if complement is directly activated by pathological α-syn or secondarily following neurodegeneration. We used the rat α-syn preformed fibril (PFF) mode, *in vitro* complement assays and human postmortem PD tissue to test whether pathological α-syn directly activates complement prior to overt neurodegeneration. The α-syn PFF model exhibits a protracted pathological time course and distinct temporal separation between peak α-syn aggregation and nigrostriatal degeneration; thus we quantified complement expression, activation, and regulation during the aggregation phase. Synucleinopathy induced complement activation prior to nigrostriatal degeneration, including upregulation of components of both the classical (*C1qa, C1r, C4b*) and alternative (*Cfd, Cfb*) pathways, the anaphylatoxin (*C3aR, C5aR*) and phagocytic (*CR3*) complement receptors, and activation of complement C3. During early synucleinopathy, microglia upregulated C3 which significantly correlated with synucleinopathy burden across several brain regions, including the substantia nigra pars compacta (SNc) and cortex. Concurrently, complement regulatory proteins, including CD55, CD59, neuronal pentraxin-1 (Nptx1), and the neuronal pentraxin receptor were downregulated in the synucleinopathy-affected SNc. Importantly, increased levels of C1q and iC3b along with downregulation of CD55 and NPTX1 were also observed in human postmortem PD SNc, supporting the translational relevance of our findings. Mechanistically, we demonstrate that aggregated, but not monomeric, α-syn directly binds C1q and activates the complement cascade in a C1q-dpendent manner. These data provide the first *in vivo* evidence that synucleinopathy triggers complement activation and dysregulation prior to neurodegeneration.

## Background

Parkinson’s disease (PD) is a progressive neurodegenerative disorder associated with clinical symptoms of parkinsonism (bradykinesia, resting tremor and postural instability) and a variable constellation of non-motor symptoms. Neuropathologically, PD is characterized by the degeneration of nigrostriatal dopamine neurons and accumulation of aggregated forms of the protein α-synuclein (α-syn)[16, 96]. Mutations and multiplications in the *SNCA* gene also cause rare familial forms of PD [4, 11, 23, 35, 38, 49, 53, 58, 74, 83, 94, 121], while SNPs in the SNCA gene or its regulatory elements increase the risk of developing PD [21, 87, 93]. Collectively these data strongly implicate α-syn pathology in PD pathogenesis, yet the exact mechanism(s) by which pathological α-syn drives neurodegeneration remain incompletely defined. However, a growing body of data suggests neuroinflammation initiated by synucleinopathy may contribute to PD pathogenesis.

Neuroinflammation is now widely recognized as a central feature of PD pathology, involving chronic and systemic activation of both innate and adaptive immunity (reviewed in [47]. Importantly, indices of immune activation are present in prodromal and *de novo* PD patients and correlate with the degree of clinical symptomology and/or the time of symptom onset [20, 22, 43, 55, 60, 72, 91, 97, 99, 122, 124], suggesting immune activation is an active participant in disease progression. As such, deciphering mechanisms of early-stage immune activation in PD may shed light on disease pathogenesis.

A growing body of data identifies synucleinopathy as a primary driver of the immune activation in PD. In the PD brain, activated microglia spatially and temporally correlate with Lewy pathology [14, 39], while microglia and monocyte activation is a characteristic feature of α-syn based models of PD [18, 30, 32, 85, 90, 107, 115]. Pathological α-syn also elicits an adaptive immune response, as T-cells from PD patients recognize α-syn peptides, autoantibodies targeting α-syn are found in PD patients, and Lewy bodies in the PD brain are coated with IgG [71, 92, 102]. Finally, immune activation is also a characteristic feature of other synucleinopathies (such as Multiple System Atrophy (MSA) and Dementia with Lewy Bodies (DLB)) further supporting the immunogenicity of pathological α-syn [1–3, 13, 54, 56, 103, 104, 120]. Due to the large body of data linking pathological α-syn to neuroinflammation we sought to identify early-stage immune mediators that respond directly to synucleinopathy. The complement system is a division of the innate immune system that continuously surveils the body and acts as a first responder to a wide range of potentially immunogenic stimuli, including pathogens and cellular damage. As such, we hypothesized that the complement system may act as a fundamental early-stage component of the neuroinflammatory cascade that drives immune activation in response to pathological α-syn.

The complement system consists of ∼50 proteins that circulate through the body as inactive precursors (**Fig. 1**). Activation of the complement system is initiated *via* one of three pathways (**Fig. 1a**): The classical, lectin, and alternative pathways. The different complement activating pathways are triggered by unique immunogenic stimuli; However, all pathways result in the sequential proteolytic cleavage of complement proteins that ultimately converge on the cleavage-mediated activation of complement component 3 (C3; **Fig. 1b**). Activation of C3 initiates the main effector responses of the complement cascade (**Fig. 1c**), including 1) the deposition of opsonins (C3b and iC3b) on targets to tag them for phagocytic clearance, 2) production of the anaphylatoxins, C3a and C5a, that recruit and activate immune cells, and 3) the lytic destruction of target cells following insertion of the membrane attack complex (MAC) in target membranes. Complement proteins are always present in tissue and can be rapidly activated and amplified, making the complement system uniquely situated to act as a first responder to immunogenic stimuli. Further, the complement system recognizes a wide range of immunogenic stimuli and coordinates inflammation following activation. As such, we predicted the complement system acts as a first responder to synucleinopathy.

**Fig. 1.**
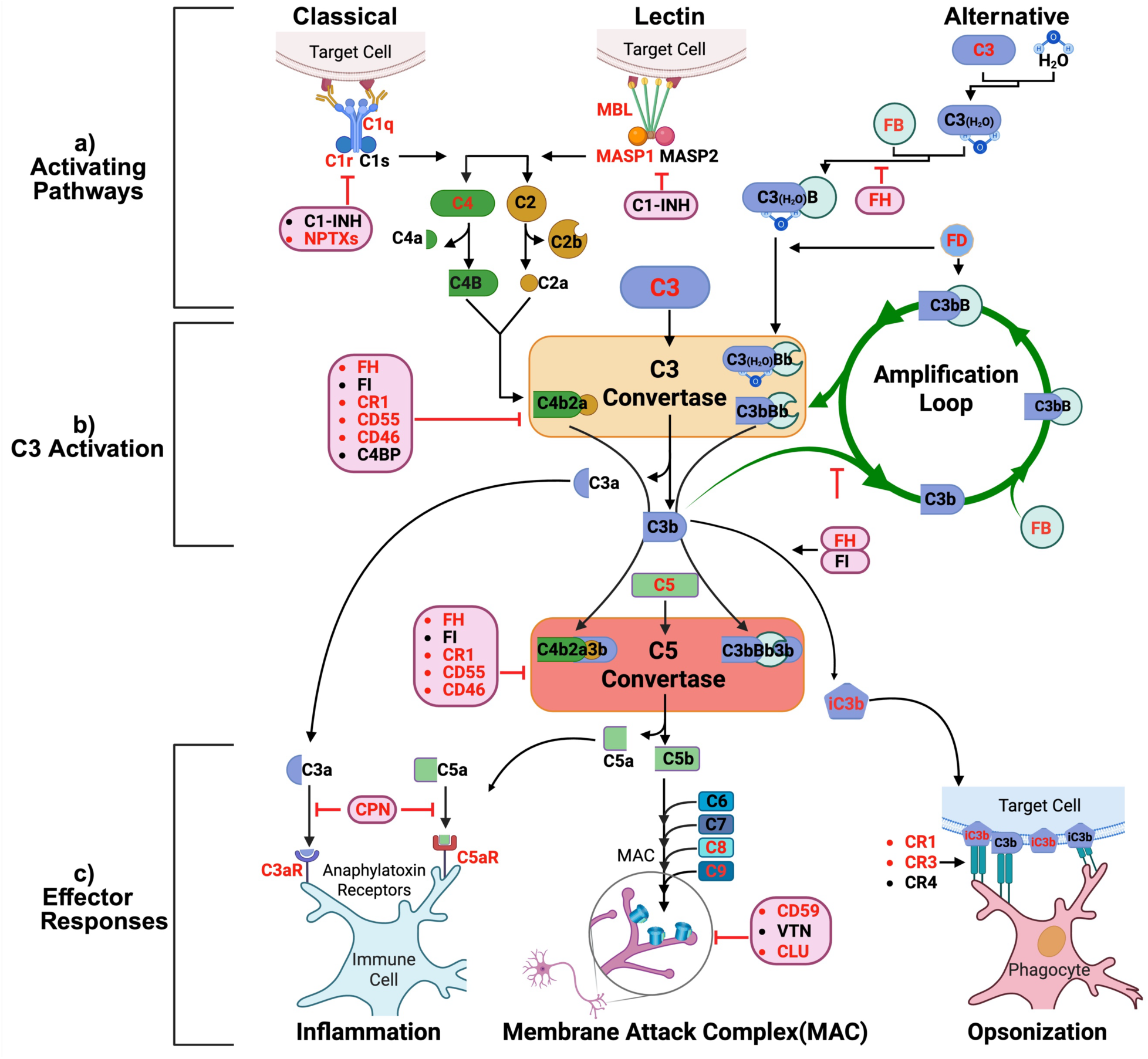
The Complement Cascade. The complement system is an arm of innate immunity, comprising ∼50 proteins present throughout 8ssues and ac8vated by proteoly8c cleavage. **a)** Complement ac8va8on occurs *via* one of three pathways: classical, lec8n, and alterna8ve. The classical pathway is ini8ated when C1q binds targets, ac8va8ng C1r and C1s, which cleave C4 and C2 to form the C3 convertase, C4b2a. The lec8n pathway is triggered by mannose-binding lec8n (MBL), collec8ns, or ficolins recognizing microbial- or altered self-carbohydrates, ac8va8ng MASP-1 and MASP-2 to generate the C3 convertase, C4b2a. The alterna8ve pathway is cons8tu8vely ac8ve through spontaneous C3 hydrolysis, forming the free C3 convertase C3(H2O)Bb, and is amplified on surfaces to form the alterna8ve pathways C3 convertase, C3bBb, enhancing complement ac8va8on. **b)** The central event in the cascade is cleavage of C3 by the C3 convertases to form C3a and C3b, which ini8ate the major effector responses of the system. **c) Opsoniza9on/Phagocytosis**: C3b and iC3b deposited on target surfaces promote phagocytosis *via* CR1, CR3, and CR4. **Inflammatory Signaling**: The anaphylatoxin pep8des, C3a and C5a, mediate inflammatory signaling through C3aR and C5aR, driving chemotaxis, cytokine produc8on, and ac8va8on of immune cells. **Membrane ACack Complex (MAC)**: C5b ini8ates the terminal pathway by recrui8ng C6–C9 to form the MAC, lysing target cells. Complement ac8vity is 8ghtly regulated at mul8ple levels (shown in pink boxes): C1 inhibitor (C1-INH) and neuronal pentraxins (NPTXs) restrain classical ini8a8on, C4 binding protein (C4BP), Factor H (FH), and FI regulate C3 convertases and alterna8ve pathway amplifica8on; carboxypep8dase-N (CPN) inac8vates anaphylatoxins; the membrane bound regulators CR1, CD46, CD55, and CD59 prevent deposi8on of ac8vated complement on self-cells; clusterin (CLU) and vitronec8n (VTN) prevent MAC assembly and inser8on. Representa8ve complement components (shown in red text) were quan8fied in α-synuclein pre-formed fibril–injected rats and postmortem PD brains.

Beyond its canonical immune functions in the periphery, the complement system is present in the CNS, where it performs various physiological functions, including coordinating synaptic pruning[98]. However, accumulating data document aberrant complement activation in neurodegenerative diseases, including PD. For instance, complement expression is markedly increased in the substantia nigra pars compacta (SNc) and caudate of PD patients[42, 66].

Additionally, multiple complement proteins are altered in the CSF and blood of PD patients, with some changes correlating with clinical symptoms and disease status[25, 26, 29, 48, 68, 69, 109, 116]. Notably, alterations in specific plasma complement proteins, including C3, predict PD up to 7 years prior to motor symptom onset, supporting the hypothesis that complement activation is an early and potentially causal contributor to disease pathogenesis in PD [29].

Like immune activation in PD, complement activation appears to target brain regions affected by synucleinopathy. In the PD brain, neuromelanin+ neurons and Lewy bodies label with activated complement proteins, including opsonins and components of the MAC [15, 62, 118]. Lewy bodies in the brains of patients with DLB also label with complement[40, 108], and complement proteins are altered in the CSF of patients with MSA [113]. Complement labeled neurons and Lewy bodies in the brains of synucleinopathy patients are surrounded by microglia expressing complement receptor 3 (CR3) and CR4 [15, 108], suggesting complement coordinates the phagocytic clearance of pathological α-syn by microglia. Consistent with this hypothesis, complement activation is also observed in preclinical α-syn-based models of PD[12, 64, 75, 100, 119, 126]. Collectively, these data strongly imply pathological α-syn drives complement activation in PD and other synucleinopathies. However, a key unresolved question is whether the complement response to synucleinopathy occurs following cell death- or prior to cell death, where it may drive neurodegeneration.

To date, most studies documenting complement activation in PD or preclinical models of PD report complement activation after significant cell death has already occurred. For instance, at the time of PD motor symptom onset, approximately 50-60% of nigral dopamine neurons have degenerated[51]. As such, complement activation documented in symptomatic PD patients could be the result of synucleinopathy, or degenerating neurons. Similarly, most studies investigating complement activation in models of synucleinopathy report complement activation at a time when significant degeneration has occurred[12, 64, 126].

In contrast, we recently reported that SNc glia rapidly upregulate expression of complement genes months prior to degeneration in the rat α-syn preformed fibril (PFF) model of synucleinopathy [77, 100]. We extensively characterized the time course of events following intra-striatal injection of α-syn PFFs in rats and identified two distinct pathological phases in the SNc [17, 18, 76, 79, 101]. The first phase or “aggregation phase” occurs 1-2 months post-injection and is characterized by peak accumulation of α-syn aggregates in nigral neurons.

These “Lewy body-like” aggregates are composed of endogenous α-syn phosphorylated at serine 129 (pSyn) and display many core features of human Lewy bodies, including proteinase-K resistance, thioflavin-S positivity and labeling with p62 and ubiquitin[18, 79]. The second phase, or “degenerative phase” occurs 4-6 months post-injection and is characterized by the progressive degeneration of nigrostriatal dopamine neurons[17, 76]. Importantly, we observe significant upregulation of complement *C3* and *C1q* in SNc glia during the aggregation phase, 2-4 months prior to overt neurodegeneration[100]. These results suggest synucleinopathy activates complement prior to degeneration. Thus, the purpose of the current study is to provide an in-depth investigation of complement activation during the early stages of synucleinopathy, and determine if pathological α-syn directly activates complement prior to overt neurodegeneration.

To accomplish this, we leveraged the protracted time course of pathology in the rat α-syn PFF model to quantify complement expression and activation during the aggregation phase (targets shown in red text in **Fig. 1**). We report that induction of synucleinopathy results in robust complement activation prior to overt neurodegeneration, including 1) upregulation of genes in both the classical and alternative complement pathways, 2) activation of C3 that significantly correlates with synucleinopathy burden in the SNc and cortex (Cx) of PFF injected rats, and 3) a significant downregulation of specific complement regulators, including CD55, CD59, neuronal pentraxin 1 (Nptx1) and the neuronal pentraxin receptor (Nptxr) in the SNc.

Further, we validated the downregulation of CD55 and Nptx1 in human postmortem PD SNc tissue and demonstrate that aggregated α-syn, but not monomeric α-syn, directly binds C1q and activate the classical complement system in a C1q-dependent manner. These results demonstrate robust complement activation and dysregulation during the early stages of synucleinopathy (i.e. prior to cell death), confirming the ability of pathological α-syn to directly activate the complement system. These findings support the hypothesis that early stage complement activation in response to pathological α-syn may drive immune activation and degeneration in PD.

## Materials and Methods

### Animals

Three-month old, male and female Fischer 344 rats (CDF, strain code 002; n=8/sex/group) were purchased from Charles River Laboratories and housed 2-3 per cage at the Michigan State University Grand Rapids Research Center, which is fully approved through the Association for Assessment and Accreditation of Laboratory Animal Care (AAALAC). Rats were housed in a room with a 12-hour light/dark cycle and provided food and water *ad libitum.* All procedures were performed in accordance with federal, state and institutional guidelines approved by the Michigan State University Institutional Animal Care and Use Committee (IACUC).

### Postmortem Human Tissue

Age matched, fresh frozen and formalin fixed paraffin embedded postmortem midbrain tissue from controls and individuals with a neuropathological diagnosis of PD was obtained from the Michigan Brain Bank (See **supplemental table 1 online resource** for detailed neuropathological and demographic data). Neuropathologic diagnoses were obtained from de-identified autopsy reports generated by board-certified neuropathologists. Detailed pathologic features were extracted from the reports, including descriptions of substantia nigra degeneration, Lewy body pathology, and co-occurring neuropathologies. Lewy pathology distribution was categorized using standard criteria (brainstem-predominant, limbic/transitional, or diffuse neocortical) when supported by the report text. Alzheimer disease neuropathologic change was classified according to NIA-AA guidelines when Braak stage and/or CERAD plaque scores were available. Features not explicitly reported were left unassigned. For biochemical analysis of postmortem tissue, fresh frozen midbrain tissue was placed on a cooling stage and viewed under a dissecting microscope. Tissue punches from the SNc were obtained with a round, 2mm diameter tissue punch inserted 2mm (2mm x 2mm tissue punch) and frozen at −80C until further processing.

### Production of Recombinant α-syn monomers and PFFs

To produce recombinant mouse α-syn, a plasmid encoding wild type (WT) mouse α-syn (PRK172, kindly gifted by Luk lab) was transformed into BL21 (DE3) RIL *E. coli* (ThermoFisher C600003) and cultured overnight at 37°C in Terrific Broth (TB) containing carbenicillin (100μg/mL) with shaking at 250 RPM. For recombinant human α-syn, a plasmid encoding WT human α-syn (Addgene Plasmid #36046) was transformed into One Shot BL21 (DE3) Star *E. coli* (ThermoFisher, C601003) and cultured at 37°C in TB containing carbenicillin (100μg/mL) with shaking at 250 RPM. When cultures reached an optical density of 0.6 at A600, protein expression was induced by adding isopropyl β-D-1-thiogalactopyranoside (IPTG) to a final concentration of 1 mM, and cultures were grown an additional three hours.

Cultures were centrifuged at 5,300 x g for 20 minutes at 4°C, supernatant decanted, and pellets collected. Pellets were resuspended in ice-cold High Salt Buffer (750 mM NaCl, 10 mM Tris, 1 mM EDTA, 1 mM PMSF, with protease inhibitor cocktail tablet (Sigma P8340)). Pellets were homogenized by three rounds of sonication using a Qsonica Q125 Sonicator connected to a 20mm horn set at amplitude 70%, with 30 second pulses and 30 second rest between pulses. The homogenate was boiled for 20 minutes, chilled on ice, and centrifuged at 5,300 x g for 20 minutes at 4°C. Supernatant was dialyzed overnight at 4°C in Buffer A (10 mM Tris pH 7.6, 1 mM EDTA). Retentate was then centrifuged 5,300 x g for 20 minutes at 4°C and filtered with a 0.45 µm PVDF filter.

α-syn was purified with two rounds of ion exchange FPLC (Cytiva AKTA Pure 25; Unicorn 7.6). Protein was loaded onto a HiPrep Q FF 16/10 column and eluted against a linear gradient of 7 column volumes of Buffer B (10 mM Tris pH 7.6, 1 mM EDTA, 1 M NaCl). Collected fractions were run on an AnyKD Criterion TGX gel (BioRad, 5671124) and stained with Coomassie blue (ThermoFisher 20279). Fractions containing a band the size of α-syn (∼14 kDa) were dialyzed overnight at 4°C in Buffer A. Protein was then loaded onto a HiTrap Q HP column and eluted against a linear gradient of 9 column volumes of Buffer B (10 mM Tris pH 7.6, 1 mM EDTA, 1 M NaCl). Collected fractions were run on a AnyKD Criterion TGX gel as before and fractions with a single band at ∼14 kDa were dialyzed overnight at 4°C in Buffer A. Endotoxin removal was performed with Pierce High-Capacity Endotoxin Removal Spin Columns (ThermoFisher 88274) per manufacturer instructions, then dialyzed overnight at 4°C in 10 mM Tris pH 7.6 containing 50 mM NaCl. Retentate was centrifuged at 10,000 x g for 30 minutes at 4°C, protein content quantified by A280 on a NanoDrop 8000 spectrophotometer (ThermoFisher) and diluted to a concentration of 5 mg/mL. A portion of purified α-syn was flash-frozen in ethanol supercooled on dry ice and stored at −80°C as α-syn monomer control. The remaining recombinant α-syn was used to generate PFFs. For fibrilization, 300 µL aliquots of recombinant α-syn monomers (5mg/mL) were shaken at 1,000 RPM for 7 days at 37°C[78]. Fibrils were aliquoted, flash-frozen in ethanol supercooled on dry ice chilled and stored at −80°C. Endotoxin contamination was tested using the *Limulus* amebocyte lysate assay and all PFF preparations contained <0.5 endotoxin units/mg of total protein. Both monomer and fibrils underwent *in vitro* testing (described below) and *in vivo* testing in rodents to ensure seeding efficacy before use in experiments.

### Recombinant α-syn PFF In Vitro Validation and Sonication

Recombinant α-syn PFF amyloid structure, oligomerization and length were confirmed *in vitro* using Thioflavin-T (ThT) fluorescence, a sedimentation assay and transmission electron microscopy, respectively (**Supplemental Fig. 1 online resource**). Amyloid structures in recombinant full-length PFFs were confirmed using a ThT assay as described [78]. Briefly, α-syn monomers and PFFs were diluted in Dulbecco’s phosphate buffered saline (dPBS) to a concentration of 0.1mg/mL and incubated with ThT in glycine buffer (25μM ThT, 100 mM glycine, 1% Triton X-100, pH 8.5) for 1 hour at room temperature (RT). ThT fluorescence was quantified using a BioTek Synergy H1 plate reader (Agilent) with excitation of 450 nm and emission of 510 nm (**Supplemental Fig. 1a-b online resource**). Aggregated α-syn in PFF preparations was confirmed using a sedimentation assay as described [78]. Briefly, α-syn monomers and PFFs were diluted in dPBS to a concentration of 0.5mg/mL and centrifuged at 10,000 x g for 30 minutes at RT, after which supernatants were transferred to a new tube.

Protein in the supernatant and pellet fraction were separated by SDS-PAGE using AnyKD Criterion TGX gel (BioRad, 5671124). Gels were then stained with Coomassie blue (ThermoFisher 20279) and α-syn resolving in the supernatant and pellet fractions from monomer and PFF preparations were validated (**Supplemental Fig. 1c-d online resource**).

Prior to stereotactic injection, PFFs were thawed to RT and diluted to 4mg/mL in sterile dPBS and sonicated with a Qsonica Q700 Cuphorn sonicator connected to a thermocube cooler set to 15°C. PFFs were sonicated at amplitude 30 with cycles of 3 seconds of sonication followed by a 2 second pause, for a total of 15 minutes of sonication. Samples were removed and gently triturated to mix the solution and then sonicated at amplitude 30 for another 7.5 minutes, again with cycles of 3 seconds of sonication followed by a 2 second pause at 15°C.

Optimal size of sonicated mouse α-syn PFFs was confirmed using transmission electron microscopy (EM; **Supplemental Fig. 1e-g online resource**). For EM of PFFs, Formvar/carbon-coated copper grids (EMSDIASUM, FCF300-Cu) were washed twice with ddH2O, then sonicated fibrils were diluted 1:50 in sterile dPBS and 10ul of diluted PFFs were adsorbed on grids for 1 minute, followed by a 1-minute incubation in 10µl of aqueous 2% uranyl acetate (Electron Microscopy Sciences 22400). Grids were allowed to dry fully before being imaged with a JEOL JEM-1400+ transmission electron microscope. Fibril lengths were measured from > 500 individual fibrils using ImageJ [89] to ensure samples contained PFFs with fibril length ≤ 50 nm, which is required for successful cellular uptake and seeding of endogenous α-syn[106]. The mean length of sonicated PFFs was 36 ± 14 nm (**Supplemental Fig. 1e-g online resource**).

### Stereotaxic Injections

Intra-striatal injection of sonicated α-syn PFFs or dPBS vehicle control were conducted as previously described[76, 100]. Rats were anesthetized with isoflurane (5% induction and 1.5% maintenance) and received unilateral intra-striatal injections to the left hemisphere (2 injections of 2 µl each). Site 1: AP +1.0, ML +2.0, DV −4.0; Site 2: AP +0.1, ML +4.2, DV −5.0. All AP and ML coordinates are relative to Bregma, while DV coordinates are relative to dura. PFFs (4 mg/mL; 2 x 2μl injection; 16 ug total) or an equal volume of dPBS vehicle were injected at a rate of 0.5 µl/minutes with a pulled glass capillary tube attached to a 10 µl Hamilton syringe. After each injection the needle was left in place for 1 minute, retracted 0.5 mm then left for another 2 minutes before fully retracting to prevent reflux. All animals received analgesic (1.2 mg/kg of sustained release buprenorphine; Ethiqa XR) and were monitored daily for 4 days post-surgery to ensure full recovery.

### Euthanasia and Tissue Preparation

Animals were euthanized at 2-months (60 days) post-injection using an injection of pentobarbital (30mg/kg i.p.; Euthanasia-III Solution, Med-Pharmex Incorporated) and perfused intracardially with heparinized (10,000 units/L) 0.9% saline. For ddPCR and protein biochemistry, brains were removed and immediately flash frozen in 2-methylbutane supercooled on dry ice for 10-20 seconds. Frozen brains were stored at −80IZ°C until further processing. Regions of interest (SN and ST) were microdissected from frozen brains using a modified version of the Palkovits punch technique[73]. Briefly, frozen brains were mounted in OTC (Fisher Healthcare Tissue-Plus OCT) and sectioned on a 305S cryostat (Leica Biosystems) set at −15IZ°C to the level of the rostral ST (∼1.96mm posterior from bregma). The ST was microdissected using a 2mm round biopsy punch, inserted to a depth of 2mm (2mm x 2mm tissue punch). Brains were further sectioned to the level of the rostral SN (∼ 4.5mm posterior from bregma), and the SN (including the SNc) was micro-dissected using a custom biopsy punch (0.5mm x 1.5mm), inserted to a depth of 2mm (∼1mm x 2mm tissue punch). Tissue was collected in DNase/RNase free microcentrifuge tubes containing 100µl TRIzol reagent (Invitrogen 26696026), homogenized with a disposable pestle, the volume of TRIzol was brought to 1mL, triturated until homogenous, and samples were frozen on dry ice and stored at −80IZ°C. For histology, animals were perfused with 4% paraformaldehyde following saline perfusion.

Brains were removed and post-fixed in 4% paraformaldehyde overnight, then cryopreserved in 15% sucrose in 0.1M phosphate buffer until saturated, followed by 30% sucrose in 0.1M phosphate buffer until saturated. Brains were maintained in 30% sucrose solution until sectioning. To section, brains were frozen with dry ice on the platform of a freezing stage, sliding knife microtome and sectioned coronally at 40µm into 6 series of sections spanning the entire rostral-caudal axis of the brain. Sections were stored in 30% sucrose, 30% ethylene glycol, in 0.1M phosphate buffer, pH 7.3 at 4°C.

### RNA Isolation

RNA isolation was conducted as described previously[100]. Tissue punches in TRIzol were thawed on ice, briefly centrifuged then transferred to Phasemaker tubes (ThermoFisher A33248) and incubated at RT for 5 minutes. Chloroform (200µl/sample) was added and tubes were mixed thoroughly by hand then incubated at RT for 10 minutes, followed by centrifugation at 16,000 x g at 4IZ°C for 5 minutes. Following centrifugation, the aqueous phase was transferred to a clean RNase free tube, and an equal volume of 100% ethanol was added to tubes and vortexed well. RNA was purified using an RNA clean and concentrator kit (Zymo Research, R1016) according to the manufacturer’s instructions with slight modification. Sample was added to the column, centrifuged for 1 minute at 12,000 x g, then columns were washed with 400µl of RNA wash buffer and centrifuged for 1 minute at 12,000 x g. DNA was removed by incubating column membranes with 3 Units of DNase I (Thermo Scientific EN0521) at RT for 15. Columns were centrifuged at 12,000 x g for 1 minute to remove DNase, then 400µl of RNA prep buffer was added and columns were centrifuged at 12,000 x g for 1 minute. Columns were washed with 700µl RNA wash buffer, centrifuged at 12,000 x g for 1 minute, washed with another 400µl of RNA wash buffer and centrifuged at 12,000 x g for 1 minute. A final centrifugation was done at 12,000 x g for 3 minutes and columns were allowed to air dry at RT for 5 minutes to remove residual wash buffer. DNase/RNase free water (30µl/sample) was added to the column and incubated for 5 minutes then eluted by centrifugation at 10,000 x g for 1 minute. Quality and quantity of RNA was assessed with Agilent 2100 Bioanalyzer using an Agilent RNA 6000 Pico Kit (5067-1513). RNA was diluted to 1IZng/µL with DNase/RNase free water, aliquoted and stored at −80IZ°C.

#### ddPCR

ddPCR was performed as previously described[100]. cDNA was prepared from 10ng of purified RNA using iScript Reverse Transcription Supermix (Bio-Rad, 1708841) according to manufacturer’s instructions, then diluted 1:1 with cDNA storage buffer (10IZmM Tris HCl pH 7.5, 0.1IZmM EDTA pH 8.0) and stored at −20IZ°C. TaqMan Gene Expression Probes (Applied Biosystems) were used for ddPCR. All respective probes (See **Supplemental Table 2 online resource** for probe details) were species specific and spanned exon-exon junctions whenever possible. For ddPCR reactions, a mastermix was made by combining 11μl of ddPCR supermix (BioRad 1863024) with 1ul of probe of interest conjugated to FAM fluorophore, and 0.5ul of *Rpl13* probe conjugated to VIC fluorophore. Mastermix was combined with 11μl of cDNA and 20IZµL of the cDNA-master mix reaction was added to a DG8 droplet generator cartridge (Bio-Rad, 1864008) with 70IZµL of droplet generation oil (Bio-Rad, 1863005). A QX droplet generator (Bio-Rad, 186-4002) was used to produce droplets containing the cDNA/mastermix reaction.

Droplets (40μL) were transferred to a 96-well plate (Bio-Rad, 12001925) and sealed with pierceable foil (Bio-Rad, 181-4040) by a plate sealer (Bio-Rad, 181-4000). Plates were transferred to a thermocycler (Bio-Rad, C1000), and the PCR reaction was carried out with the following settings: 10IZminutes at 95IZ°C, 39 cycles (30IZs at 94IZ°C, 1IZminute at 60IZ°C), 10IZminutes at 98IZ°C, hold at 12IZ°C (constant lid temperature of 105IZ°C). Plates were then transferred to the QX200 droplet reader (Bio-Rad, 1864003), and results analyzed with QuantaSoft software. For all samples, the gene of interest was normalized to the reference gene, *Rpl13*, and final data was expressed as fold change from PBS controls.

### SDS-PAGE and Western Blotting

Tissue punches from the rat ST and SN, and human SNc, were homogenized in 10 volumes (w/v) of RIPA buffer (50 mM Tris pH 7.4, 150 mM NaCl, 1% NP-40, 0.5% sodium deoxycholate, 1% sodium dodecyl sulfate (SDS), 1mM Ethylenediaminetetraacetic acid (EDTA)) containing protease (10 μg/mL pepstatin, leupeptin, bestatin and aprotinin, and 1 mM Phenylmethylsulfonyl fluoride (PMSF)) and phosphatase inhibitors (1 mM tetra-sodium pyrophosphate decahydrate, 10 mM beta-glycerophosphate disodium salt pentahydrate, 1 mM sodium orthovanadate, 10 mM sodium fluoride) by sonication using 10 short pulses at power 2 (Misonix XL-200). Crude lysates were centrifuged at 13,000 x g for 20 minutes at 4° C and the supernatant containing soluble protein was saved. Protein content was determined using a Pierce rapid gold BCA kit (Thermo, A53225) according to the manufacturer’s instructions. To test the specificity of the phospho-serine 129 α-syn antibody used for biochemical quantification of pSyn, lysates were incubated overnight at 37°C with and without FastAP Thermosensitive Alkaline Phosphatase (7.5U; ThermoFisher EF0651). For all samples, 25-60 μg of protein was mixed with 6x Laemmli sample buffer and heated to 98°C for 5 minutes. Proteins were separated by SDS-PAGE using AnyKD Criterion TGX gels (BioRad, 5671124) and transferred to BioTrace nitrocellulose membranes (VWR, 27376-991). Membranes were blocked in tris buffered saline (TBS; 0.15M sodium chloride, 0.05M Tris, pH7.6) containing 2% non-fat dry milk (Sanalac) for 1 hour prior to overnight incubation at 4°C in primary antibodies. Primary antibodies and the concentrations used are listed in **Supplemental 3 online resource**.

Membranes were then washed 3 x 5 minutes in tris-buffered saline (TBS) containing 0.1% tween-20 (TBS-T). For immunoblotting of rat C3 only, membranes were incubated in a goat anti-rabbit horseradish peroxidase conjugated secondary antibody (Jackson ImmunoResearch 111-035-144; RRID:AB_2307391) diluted at 1:5,000 for 2 hours at RT, then developed with an enhanced chemiluminescent substrate (BioRad 1705062) according to manufacturer’s instructions. Rat C3 membranes were washed 3 x 5 minutes in TBS before being imaged on a Biorad Chemidoc Imager. For all other target proteins, membranes were incubated in either goat anti-mouse IRDye 800CW (LiCor 926-32210; RRID: AB_621842), goat anti-rabbit IRDye 680RD (LiCor 926-68071; RRID: AB_2313606), or donkey anti-goat IRDye 680RD (LiCor 926-68074; RRID: AB_10956736) diluted at 1:20,000 for 2 hours at RT. Membranes were washed 3 x 5 minutes in TBS-T and then imaged on a LiCor Odyssey near-infrared imaging system with ImageStudio software (v5.2.5, LiCor Biosciences).

### Sandwich Enzyme Linked Immunosorbent Assay (sELISA)

A sELISA was developed to quantify levels of human iC3b in tissue lysates generated from the SNc of control and PD brains (see SDS-Page and Western Blotting for a description of lysate preparation). Levels of iC3b were measured using a monoclonal antibody raised against a neoepitope in human iC3b (Quidel A209) as a capture antibody and a polyclonal anti-human C3 antisera (Quidel A304) as a detector antibody (**Supplemental Fig. 2 online resource**). The iC3b capture antibody was diluted to 2ng/μl in borate saline (100IZmM boric acid, 25IZmM sodium tetraborate decahydrate, 75IZmM NaCl, 250μM thimerosal) and 50μl was added to wells of high binding microtiter plates (Corning 3590) and incubated overnight at 4°C. Wells were rinsed twice with 200μl of ELISA wash buffer (100IZmM boric acid, 25IZmM sodium tetraborate decahydrate, 75IZmM NaCl, 250 μM thimerosal, 0.4% bovine serum albumin, and 0.1% tween-20) and blocked with 200μl ELISA wash buffer containing 5% nonfat dried milk (Sanalac) for 1IZhour at RT. Wells were then washed twice with 200μl ELISA wash buffer. A standard curve of purified human iC3b protein (Complement Tech, A115) was generated by performing 1:3 dilutions of iC3b in TBS, with the standard curve ranging from 200nM to 0.001nM, and 50μl/well were added to plates in triplicate. Human brain lysate were diluted to a final concentration of 1μg/μl, and 50μl/well was added to plates in duplicate. Wells containing TBS and RIPA lysis buffer blanks were included to control for non-specific signal generated from buffers. Standards, samples and blanks were incubated in wells for 2 hours at RT. Wells were washed four times with 200μl of ELISA wash buffer, after which the anti-human C3 detector antibody was diluted 1:500 in ELISA wash buffer containing 5% non-fat dry milk, and 50μl was added to each well and incubated for 2 hours at RT. Wells were washed four times with 200μl ELISA wash buffer and then incubated with 50μl/well of a donkey-anti goat secondary antibody conjugated to horseradish peroxidase (Jackson ImmunoResearch 705-035-003; RRID: AB_2340390), diluted 1:5,000 in ELISA wash buffer containing 5% non-fat dry milk. Wells were washed 4 times with 200μl ELISA wash buffer and bound antibody was detected with 50μL of 3,3’,5,5’-tetramethylbenzadine (TMB; Sigma Aldrich T0440) substrate for 2-15 minutes. The reaction was stopped with 50μL of 3.6% H_2_SO4 and absorbance at 450nm (A450) was quantified with a Spectramax plus plate reader (Molecular Devices 18780). The average A450 of blank wells was subtracted from all wells to account for non-specific background signal, and the average from standard triplicates and sample duplicates was calculated. Absorbance (A) is not linear [i.e., A = Log10(1/transmittance)], thus, the absorbance data were converted to percent absorbed light (a linear scale) using the following equation percent light absorbed = (1 – 10−x)∗100, where x is absorbance. The standard curve was Log_10_ transformed and best fit to a sigmoidal curve, thereby providing a standard curve of known amounts of iC3b protein. Samples were plotted onto the standard curve to ensure all sample values were within the linear range of the assay (**Supplemental Fig. 2 online resource**), and the quantity of iC3b protein (ng) in each human sample was then interpolated from the iC3b standard curve and converted to a concentration expressed in ng/μl. Graphs and concentrations were produced using GraphPad Prism 11 software.

### Immunohistochemistry

Immunohistochemistry was performed on both free-floating rat tissue and paraffin embedded human tissue. For paraffin embedded tissue, tissue was de-paraffinized by performing 2 x 5 minute incubations in xylenes, 2 x 3 minute incubations in 100% ethanol, 2 x 1 minute incubations in 95% ethanol, a 1 x 1 minute incubation in 70% ethanol, a 1 x 1 minute incubation in 50% ethanol followed by a ∼15 second dip in ddH2O. Tissue was then incubated in 1x IHC Select Citrate Buffer (Millipore 21545) at 95°C for 10 minute to expose epitopes, and then cooled to RT.

All tissue (both free floating and paraffin embedded) was then washed 6 x 10 minutes in TBS containing 0.5% triton X-100 (TBS-Tx). Endogenous peroxidase activity was quenched by incubating tissue in 3% hydrogen peroxide diluted in TBS-Tx for 1 hour, followed by 6 x 10 minutes washes in TBS-Tx. Tissue was blocked for 1 hour in TBS-Tx containing 10% goat serum and 2% bovine serum albumin (BSA). Tissue was then incubated overnight at RT in primary antibody diluted in TBS-Tx containing 2% goat serum. Primary antibodies and concentrations used are listed in **Supplemental table 3 online resource**. Following primary antibody incubation, tissue was washed 6 x 10 minutes in TBS-Tx. After which, the tissue was incubated in goat anti-rabbit biotinylated (Vector Laboratories, BA-1000; RRID: AB_2313606) or goat anti-mouse biotinylated (Jackson Immunoresearch115-065166; RRID AB_2338569) secondary antibodies diluted at 1:500 in TBS-Tx containing 2% goat serum for 2 hours at RT. Tissue was then washed 6 x 10 minutes washes in TBS-Tx and in incubated in ABC Elite solution (Vector Labs, PK6100) according to manufacturer’s instructions for 2 hours at RT. Tissue was washed 6 x 10 minutes in TBS-Tx, and bound antibodies were visualized using 3, 3’-diaminobenzidine (Sigma, D5637) at 0.5 mg/ml in TBS-Tx containing 0.003% hydrogen peroxide for 5-10 minutes. Tissue was then washed in TBS, mounted on microscope slides and processed through ethanol and xylenes before cover slipping with Cytoseal-60 (Thermo, 831016). Images were acquired with a Nikon Eclipse 90i microscope, a Nikon DS-Ri1 camera and Nikon Elements Software (Nikon Instruments, Melville, NY). All individual images between animals within a particular stain were acquired using identical microscope and post-processing parameters (magnification, light source intensity, exposure time and contrast). Images were prepared for publication using Adobe Photoshop (version 26.8.1) and Illustrator (version 29.6.1).

### Immunofluorescence and Quantification

Immunofluorescence staining was performed using established protocols [7, 100]. Fixed coronal sections were washed 6 x 10 minutes in TBS containing 0.5% Triton-x 100 (TBS-Tx). For C3 staining in rat tissue only (using Abcam ab200999), tissue was then incubated in 1x IHC Select Citrate Buffer (Millipore 21545) at 95°C for 10min to expose epitopes, cooled to RT, then washed 6 x 10 minutes in TBS-Tx. All tissue was blocked for 1 hr at RT in TBS-Tx containing 10% goat serum and 2% bovine serum albumin. Tissue was then incubated overnight at RT in primary antibody diluted in TBS-Tx containing 2% goat serum. Primary antibodies and the concentrations used are listed in **Supplemental Table 3 online resource**. Following primary antibody incubation tissue was washed 6 x 10 minutes in TBS-Tx, and then incubated for 2 hours at RT in secondary antibody diluted 1:500 in TBS-Tx containing 2% goat serum. Secondary antibodies used were goat anti-rabbit IgG Alexa Fluor 488 (Invitrogen, A11034; RRID: AB_2576217), goat anti-rabbit IgG Alexa Fluor 594 (Invitrogen, A11012; RRID: AB_2534079), goat anti-rabbit Alexa Fluor 647 (Invitrogen, A21245; RRID: AB_2535813), goat anti-mouse IgG2a Alexa Fluor 594 (Invitrogen, A21135; RRID: AB_2535774), goat anti-mouse IgG2a Alexa Fluor 488 (Invitrogen, A21131; RRID_AB_141618), Goat anti mouse IgG1 Alexa Fluor 488 (Invitrogen, A11029; RRID: AB_2534088), Goat anti-mouse IgG2b Alexa Fluor 488 (Invitrogen, A21141; RRID: AB_2535778), goat anti-mouse IgG2b Alexa Fluor 647 (Invitrogen, A21242; RRID: AB_2535811). Tissue was washed 6 x 10 minutes in TBS-Tx with the first of 6 washes containing 4’,6-Diamidino-2-Phenylindole, Dihydrochloride (DAPI) (1:10,000, Invitrogen, D1306: RRID: 2629482). Sections were mounted on HistoBond+ slides (VWR VistaVision, 16004-406) and cover-slipped with VECTASHIELD hardset antifade mounting medium (Vector Laboratories, H-1400-10). Microscope slides were then imaged on a Nikon Eclipse 90i fluorescence microscope or a Nikon A+ laser scanning confocal microscope and Nikon Elements Software (Nikon Instruments, Melville, NY). All individual images between animals within a particular stain were acquired using identical microscope and post-processing parameters (magnification, light source intensity, laser intensity, exposure time and contrast).

Quantification of Immunofluorescent intensity was performed using FIJI (version 2.14.0/1.54f). Briefly, images were imported into FIJI and a threshold was applied to exclude background fluorescence. Identical thresholding parameters were applied to all images between animals within a stain. A region of interest was drawn around the anatomical boundaries of the brain regions being analyzed and the “analyze particles” function was used to quantify the fluorescent intensity and percent area of positive staining within the region of interest.

Integrated density is reported as an index of fluorescent intensity. Images were prepared for publication using Photoshop (version 26.8.1) and Illustrator (version 29.6.1).

### C1q Binding Assay

A plate based-C1q binding assay was developed to quantify the ability of human C1q to bind human α-syn monomers and human α-syn PFFs. Human α-syn monomers were purified and fibrillized as described above. Purified human α-syn monomer, human α-syn PFF, human serum albumin (HSA; Sigma Aldrich A9511) or human IgG (Jackson ImmunoResearch 009-000-003; RRID:AB_2337043) proteins were diluted to a concentration of 12μM in phosphate buffered saline (PBS; 137mM NaCl, 2.7mM KCl, 10mM Na_2_hPO_4_, 1.8 mM KH_2_PO_4_) and serially diluted 1:3 from 12μM to 0.2nM in PBS. 50μL of the respective dilutions were added to wells of high binding microplates (Corning 3590) and incubated overnight at 4°C with gentle shaking. Wells were washed 3 times with 200μL PBS containing 0.05% tween-20 (PBS-T), then blocked with 200μL of 5% non-fat dry milk (Sanalac) diluted in PBS for 1 hour at RT with gentle shaking. Wells were washed 3 times with 200μL PBS-T. Purified human C1q (complement technology A099) was diluted to 2μg/mL in GVB++ buffer (complement technology B102) pre-heated to 37°C and 50μL was added to wells and incubated for 1 hour at 37°C with gentle shaking. Wells were washed 4 times with 200μL PBS-T and then incubated in 50μL of a monoclonal anti-human C1q primary antibody (1:2,000, Invitrogen MA1-40311; RRID:AB_2067275) diluted in 5% non-fat dry milk for 1 hour at RT with gentle shaking. Wells were washed 4x with 200μL PBS-T and then incubated in 50μL of goat anti-mouse horse radish peroxidase conjugated secondary antibody (1:5,000; Jackson ImmunoResearch 115-035-003; RRID:AB_100015289) diluted in 5% non-fat dry milk for 1 hour at RT with gentle shaking. Wells were washed 4x with 200μl PBS-T and then bound antibodies were visualized with 50μL of 3,3’,5,5’-tetramethylbenzadine (TMB; Sigma Aldrich T0440) substrate for 5-15 minutes. The reaction was stopped with 50μL of 3.6% H_2_SO4 and absorbance at 450nm (A450) was quantified with a Spectramax plus plate reader (Molecular Devices 18780). The average A450 of blank wells (containing only 5% non-fat dry milk) was subtracted from all wells to account for non-specific background signal. As above, the absorbance data were converted to percent absorbed light (a linear scale) using the following equation percent light absorbed = (1 – 10−x)∗100, where x is absorbance. Graphs and EC50 concentrations were produced using GraphPad Prism software.

### Complement Activation Assay

A plate based-complement activation assay was developed to quantify the ability of human α-syn monomers or human α-syn PFFs to activate the complement system. Human α-syn monomers were purified and fibrillized as described above. Purified human α-syn monomer, human α-syn PFF, human serum albumin (HSA: Sigma Aldrich A9511) or human IgG (Jackson Immunoresearch 009-000-003; RRID:AB_2337043) proteins were diluted to a concentration of 4μM in phosphate buffered saline (PBS; 137mM NaCl, 2.7mM KCl, 10mM Na_2_hPO_4_, 1.8 mM KH_2_PO_4_) and serially diluted 1:3 from 4μM to 0.067nM in PBS. Samples (50μL/well) were added to high binding 96 well microplates (Corning 3590) and incubated overnight at 4°C with gentle shaking. The following day, wells were washed 3 times with 200μL PBS containing 0.05% tween-20 (PBS-T) and then blocked in 2% bovine serum albumin (BSA; Fisher BP1600-100) diluted in PBS for 1 hour at RT. Wells were washed three times with 200μL PBS-T and then incubated in 2% normal human serum (NHS; Complement Technology, NHS) diluted in GVB++ buffer (complement technology B102) at 37°C for 1 hour with gentle shaking. Wells were then washed four times with 200μL of PBS-T and incubated in 50μL of a primary antibody specific to a neo-epitope in cleaved (activated) complement C3 (1:500; Hycult HM2257; RRID:AB_1953566) diluted in 2% BSA for 1 hour at RT with gentle shaking. Wells were washed 4x with 200μL PBS-T and then incubated in 50μL of goat anti-mouse horse radish peroxidase conjugated secondary antibody (1:5,000, Jackson Immunoresearch 115-035-003; RRID:AB_100015289) diluted in 2% BSA for 1 hour at RT with gentle shaking. Wells were washed 4x with 200μl PBS-T and bound antibodies detected with 50μL of 3,3’,5,5’-tetramethylbenzadine (TMB; Sigma Aldrich T0440) substrate for 5-15 minutes. The reaction was stopped with 50μL of 3.6% H_2_SO_4_ and absorbance at 450nm (A450) was quantified with a Spectramax plus plate reader (Molecular Devices 18780). As above, absorbance data were converted to percent absorbed light (a linear scale) using the following equation percent light absorbed = (1 – 10−x)∗100, where x is absorbance. Graphs and EC50 concentrations were produced using GraphPad Prism 11 software.

### Statistical Analysis

Statistical analysis and graphing of results were performed using GraphPad Prism 11. For ddPCR and biochemical endpoints in rat tissue, data from both sexes were compiled, and a two-way ANOVA with a Tukey’s multiple comparison test was used to compare differences between sexes and treatment. If there was no significant difference between sexes within a treatment group, sexes were combined and an unpaired t-test with Welch’s correction was used to determine differences between groups. Statistical data for all analyses that were used to determine sex differences is included in the **Supplemental Tables 4, 5 online resource**. Immunofluorescence analysis of C3, CR3 and NPTX1 was performed to localize the cell source and measure protein of targets found to be significantly altered by ddPCR and immunoblotting. As no significant sex differences were detected in these targets by ddPCR or immunoblotting (**Supplemental Table 4, 5 online resource**), immunofluorescent analysis was performed on mixed sexes. For concentration response analyses in the *in vitro* C1q binding assay and complement activation assay, data were analyzed using a two-way ANOVA with Tukey’s multiple comparison. Statistical analysis of EC50 and maximum binding/maximum activation calculated for C1q binding and complement activation assays, was performed with a one-way ANOVA and Tukey’s multiple comparison test. In all cases significance was set at p ≤ 0.05.

## Results

We aimed to determine if pathological α-syn activates the complement system prior to neurodegeneration using a rat model of synucleinopathy. Two-months post-PFF injection represents the time of peak pathological α-syn accumulation and peak glial activation, but months prior to overt degeneration of nigrostriatal dopamine neurons[18, 76, 100, 101]. A subset of animals was used to confirm successful seeding of endogenous α-syn into pSyn+ aggregates and microglial activation (as indicated by MHC-II expression) in the ipsilateral SNc. In line with previous reports [18, 76], the ipsilateral SNc of α-syn PFF injected rats contained abundant pSyn+ aggregates (**Fig. 2b**), and MHC-II+ microglia (**Fig. 2d**), while no pSyn+ aggregates or MHC-II+ microglia were observed in the SNc of PBS injected rats (**Fig. 2a, c**). Next, we quantified *C3* gene expression in the SN and ST and detected a significant increase in *C3* expression in both brain regions of PFF injected rats compared to PBS controls (**Fig. 2e-f**), validating and extending previous findings [100].

**Fig. 2.**
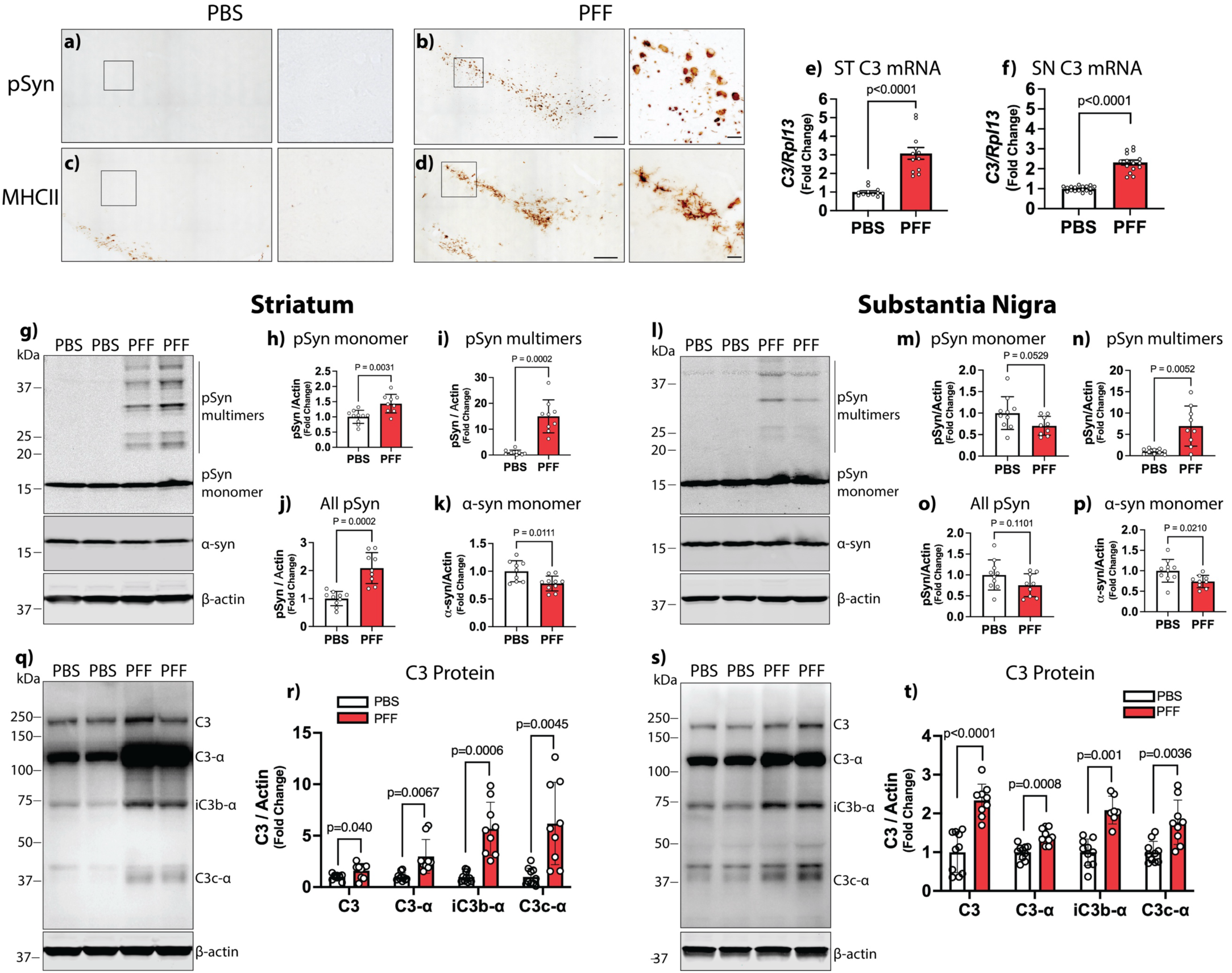
C3 Expression and Ac9va9on are Increased in a-syn PFF Treated Rats and Correlate with Levels of pSyn. Rats (n=6-8/sex/group) received intra-striatal injec8ons of α-synuclein (α-syn) preformed fibrils (PFFs) or phosphate buffer saline (PBS) and were sacrificed 2-months post-injec8on. **A-b)** Representa8ve images of Serine 129 phosphorylated α-syn (pSyn) immunostaining in the substan8a nigra (SN) of PBS (**a**) and α-syn PFF (**b**) injected rats. **C-d**) Representa8ve images of MHC-II immunostaining in the SN of PBS (**c**) and α-syn PFF (**d**) injected rats. High magnifica8on images to the right of each panel correspond to the area in the box of respec8ve low magnifica8on images. **E-F**) Droplet digital PCR (ddPCR) quan8fica8on of complement component 3 (*C3)* expression in the striatum (ST; **panel e**) and SN (**f**). Data are *C3* normalized to ribosomal potein L13 (*Rpl13*), analyzed with t-test with Welch’s correc8on. **g**) Representa8ve immunoblot of pSyn, α-syn and β-ac8n from the ST. **h**) Quan8fica8on of pSyn monomers (∼14-17 kDa), **i**) pSyn mul8mers (∼20-50 kDa), **j**) total pSyn signal (∼14-50 kDa), and **k**) α-syn monomer (∼14-17 kDa) normalized to β-ac8n in the ST. **l)** Representa8ve immunoblot of pSyn, α-syn and β-ac8n from the SN. **m**) Quan8fica8on of pSyn monomers (∼14-17 kDa), **n**) pSyn mul8mers (∼20-50 kDa), **o**) total pSyn signal (∼14-50 kDa), and **p**) α-syn monomer (∼14-17 kDa) normalized to β-ac8n in the SN. **q)** Representa8ve immunoblot of C3 and β-ac8n from the ipsilateral ST. **s**) Representa8ve immunoblot of C3 and β-ac8n from the ipsilateral SN Quan8fica8on of C3 whole molecule (∼190 kDa), C3 α-chain (∼115 kDa), iC3b α-chain (∼75 kDa), and C3c α-chain (∼34kDa) normalized to β-ac8n from the ST (**r**) and SN (**t**), analyzed with t-test with Welch’s correc8on). Scale bars in large images of (**b, d)** are 250μm and apply to large images of (**a-d)**, while scale bar in the small images of (**b, d)** are 50μm and apply to small images of (**a-d)**. All data are group means ± standard devia8on expressed a fold change from PBS group.

To determine if pathological α-syn activates the complement system prior to degeneration we performed biochemical quantification of pSyn (as a surrogate marker of pathological α-syn) and C3 in the SN and ST, but we first confirmed the specificity of the antibody used to quantify pSyn (**Supplemental Fig. 3 online resource**). There was a significant increase in the amount of pSyn monomer (∼14-17 kDa band) and pSyn multimers (∼20-50 kDa bands) in the ST of PFF injected rats (**Fig. 2g, h, i**), while levels of α-syn monomer were significantly decreased in the ST of PFF injected rats (**Fig. 2g, k**). In the SN, pSyn multimers were significantly increased (**Fig. 2l, n**), while α-syn monomers were significantly reduced (**Fig. 2l, p**).

Complement C3 sits at a functional nexus in the complement system (**Fig. 1b**) and complement proteins are activated by proteolytic cleavage. Thus, to quantify complement activation in the ST and SN of PBS and PFF injected rats, we performed immunoblotting to quantify intact C3 (as indexed by whole molecule C3 (∼190 kDa), the α-chain of intact C3 (∼115 kDa)), the α-chain of iC3b (∼68 kDa; an activated complement opsonin derived from C3) and the α-chain of C3c (∼35-40 kDa; a downstream product of C3 activation). In line with C3 gene expression (**Fig. 2e-f**), overall levels of C3 protein were significantly increased in the ST (**Fig. 2q-r)** and SN **(Fig. 2s-t)** of α-syn PFF injected rats. Further, levels of iC3b and C3c were significantly increased in the ST and SN of PFF injected rats compared to controls, confirming complement activation in response to α-syn PFF injection (**Fig. 2q-t**). Together, these data show that injection of α-syn PFFs in the ST of rats induces the formation of pSer129+ α-syn multimers in the ST and SN that corresponds with a shift of soluble α-syn monomers to insoluble α-syn aggregates, and an associated increase in C3 expression and activation in the ST and SN.

We next sought to identify the cell source upregulating C3 in α-syn PFF injected rats using immunofluorescent (IF) detection of C3, pSyn and the neuronal marker, Huc/d. No pSyn immunoreactivity and very few C3+ cells were observed in the ventral midbrain of PBS injected rats (**Fig. 3a-d, i-l**), where the majority of C3+ cells were located outside of the SNc, primarily in the substantia nigra pars reticulata (SNr) and cerebral peduncles of the ventral midbrain (**Fig. 3c**). In contrast, abundant pSyn+ inclusions were present throughout the SNc of α-syn PFF injected rats, which was associated with a dramatic increase in the number of C3+ cells in the SNc (**Fig. 3e-h, m-p**). The upregulation of C3 in the midbrain of PFF injected rats spatially overlapped with pSyn aggregates in the SNc, but increased C3+ cells were also observed in the SNr and areas adjacent to the SNc. High magnification confocal imaging of the pSyn seeded SNc shows abundant C3+ cells in direct apposition to neurons containing pSyn aggregates (**arrows in Fig. 3m-p**), suggesting C3+ cells are targeting synucleinopathy-affected neurons. C3 fluorescence intensity and the percent area of C3 staining were significantly increased in the SNc of α-syn PFF injected rats (**Fig. 3q, t**), and this was paralleled by significant increases in pSyn IF intensity and percent area staining (**Fig. 3r, u**). In α-syn PFF injected rats, C3 IF intensity was significantly correlated with pSyn IF intensity (**Fig. 3s**). Finally, we identified microglia as the primary cellular source of C3 in the SNc of PFF injected rats, as indicated by high colocalization between C3 with IBA1 (**Fig. 3w-z)** and lack of C3 colocalization with GFAP, in the brains of PFF injected rats (**Supplemental Fig. 4 online resource)**. These data are in line with the biochemical quantification of C3 and pSyn in PFF injected rats (**Fig. 2**) and, utilizing complementary endpoints, validate the association between neuronal pSyn pathology and microglial C3 upregulation during the early phases of synucleinopathy in α-syn PFF injected rats.

**Fig. 3.**
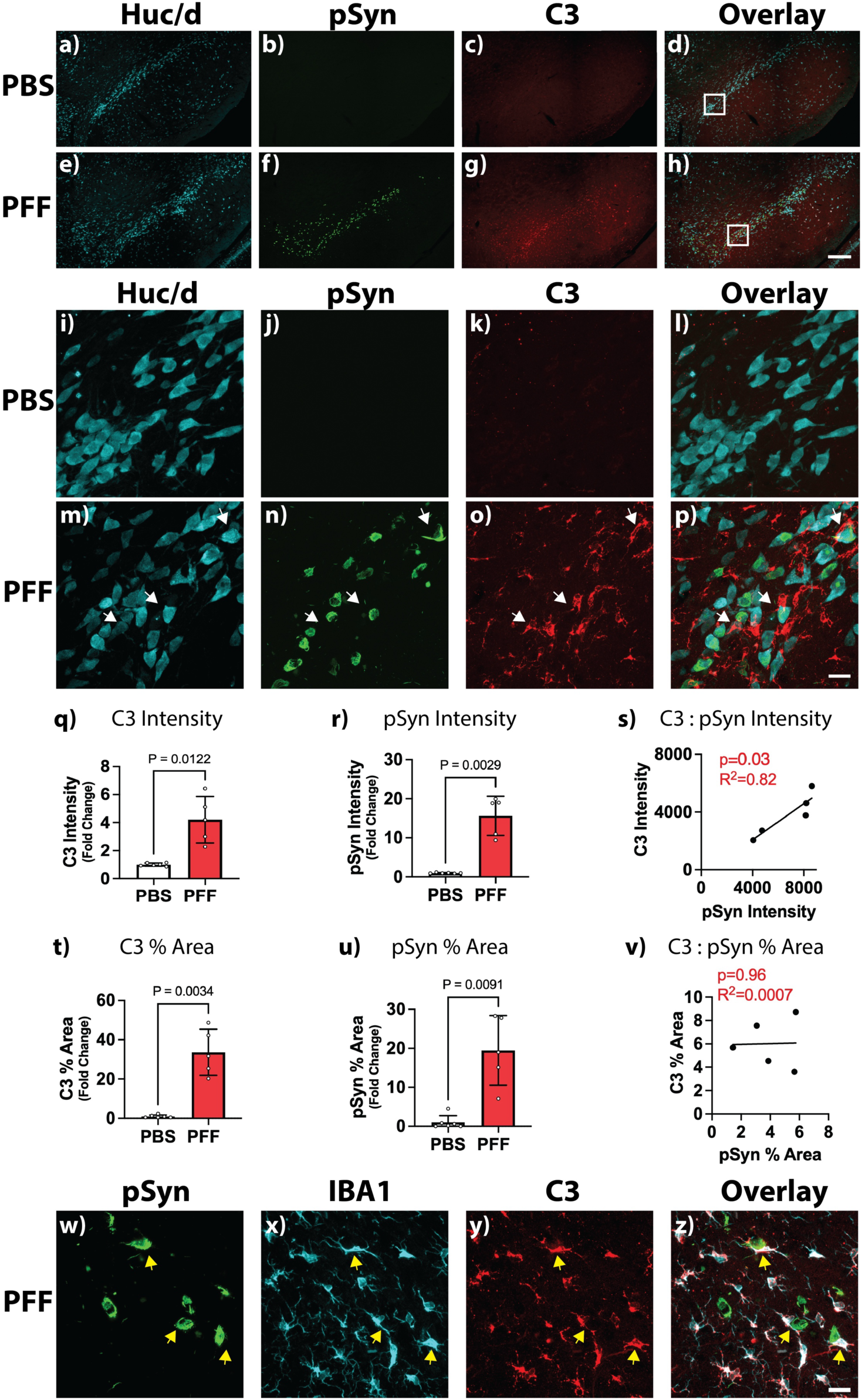
Microglia Upregulate C3 in the Substantia Nigra of a-syn PFF Injected Rats. Male and female rats (n=5-6/group) received intra-striatal injec8ons of α-synuclein (α-syn) preformed fibrils (PFFs) or phosphate buffer saline (PBS) and were sacrificed 2 months post injec8on. **a-h**) Representative low magnification immunofluorescent (IF) images of Huc/d (cyan, neuronal marker), serine 129 phosphorylated α-syn (pSyn; green) and complement component 3 (C3; red) and the overlay image in the SNc of PBS (**a-d**) and PFF (**e-h**) injected rats. **i-p**) High magnification images corresponding to box in (**d**) and (**h**), for PBS (**i-l**) and PFF injected (**m-p**) rats, respectively. **q-r**) Quantification of C3 (**q**) and pSyn (**r**) fluorescence intensity. **s**) Regression analysis between pSyn and C3 fluorescence intensity in the SNc of PFF injected rats. **t-u**) Quantification of percent area of SNc occupied by C3+ (**t**) and pSyn+ (**u**) staining. **v**) Regression analysis between pSyn and C3 percent area staining in the SNc of PFF injected rats. Data expressed as mean fold change (± standard deviation) from PBS controls, analyzed by t-test with Welch’s correction or simple linear regression analyses. **w-z**) Representative IF images of pSyn (green; panel **w**), the microglial marker, ionized calcium binding adapter molecule 1 (IBA1; cyan; panel **x**), C3 (red; panel; **y**) and the overlay image (**z**) in the SNc of PFF injected rats. Arrows in (**m-p**) and (**w-z**) indicate areas where C3+ microglia are in direct apposition of neurons containing pSyn+ aggregates. Scale bars in (**h**) is 250μm and applies to (**a-h**), scale bar in (**p**) is 50μm and applies to (**i-p**), scale bar in (**z**) is 50μm and applies to (**w-z**)

Next, we sought to determine if microglial C3 upregulation in α-syn PFF injected rats is unique to the SNc or if it also occurs in other brains regions that accumulate pathological α-syn (i.e. the cortex (Cx) and ST) [76, 79]. In α-syn PFF injected rats, pSyn immunoreactivity was significantly increased in the Cx (**Fig. 4e-h, m-p**; at the level of the claustrum, endopiriform nucleus, agranular and piriform areas) and ST (**Fig. 4aa-dd, ii-ll**). As previously described[76], pSyn immunoreactivity in the Cx appears as concentrated aggregates in neuronal somata (**Fig. 4n, p**), while pSyn immunoreactivity in the ST appears as pSyn+ neuronal processes resembling Lewy neurites, reflecting pathological α-syn in the terminals of nigrostriatal and corticostriatal neurons (**Fig. 4jj, ll**). Regardless of the brain region or morphology of pSyn aggregates, there was a significant increase in C3+ cells in both the Cx (**Fig. 4g, o**) and ST (**Fig. 4cc, kk**) of PFF injected rats. Compared to PBS injected controls, both C3 and pSyn IF intensity and percent area staining were significantly increased in the Cx (**Fig. 4q-r, t-u**) and ST (**Fig. 4mm-nn, pp-qq**) of PFF injected rats. Finally, in α-syn PFF injected rats C3 IF intensity and percent area staining significantly correlated to pSyn IF intensity and percent area staining in the Cx **(Fig. 4s, v**) but not the ST (**Fig. 4oo, rr**). These data suggest microglia upregulate C3 in response to pathological α-syn across affected brain regions.

**Fig. 4.**
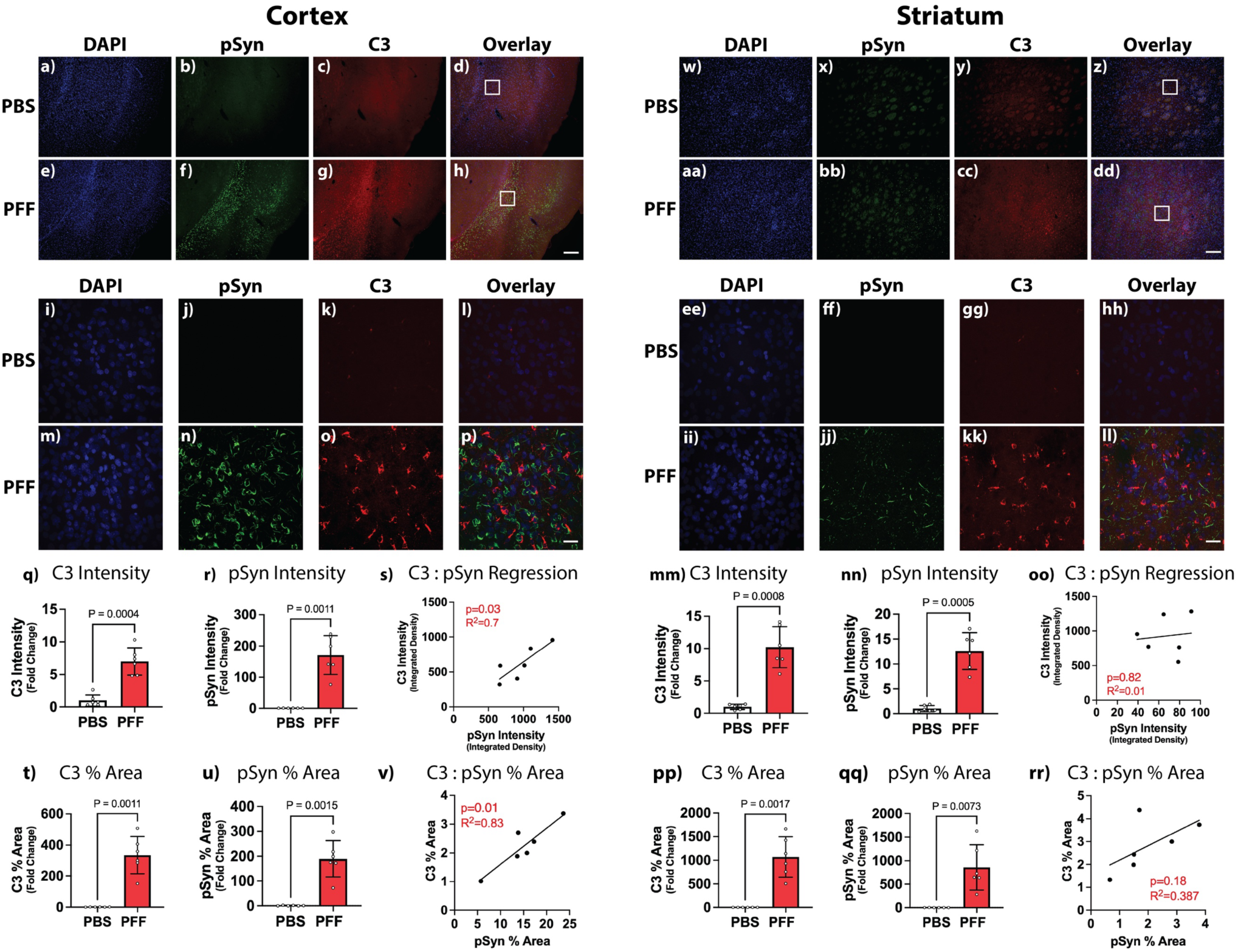
C3 Upregulation in the Striatum and Cortex of a-syn PFF Injected Rats. Male and female rats (n=5-6/group) received intra-striatal injec8ons of α-synuclein (α-syn) preformed fibrils (PFFs) or phosphate buffer saline (PBS) and sacrificed 2-months post-injec8on. **a-h**) Representative low magnification images of DAPI (blue, nuclear marker), pSyn (green, phospho-Ser129 α-syn) and C3 (red, complement 3) and the overlay in the ipsilateral cortex (Cx) of PBS (**a-d**) and α-syn PFF (**e-h**) injected rats. **i-p)** High magnification images corresponding to box in (**d**) and (**h**), for PBS (**i-l**) and PFF (**m-p**) injected rats, respectively. **q-r**) Quantification of C3 (**q**) and pSyn (**r**) fluorescence intensity in the Cx. **s**) Linear regression between pSyn and C3 fluorescence intensity in the Cx of PFF injected rats. **t-u)** Quantification of percent area of Cx occupied by C3+ (**t**) and pSyn+ (**u**) staining. **v**) Linear regression between pSyn and C3 percent area staining in the Cx of PFF injected rats. **w-dd**) Representative low magnification images of DAPI (blue), pSyn (green) and C3 (red) and overlay image in the ipsilateral striatum (ST) of PBS (**w-z**) and PFF (**aa-dd**) injected rats. **ee-ll)** High magnification images corresponding to box in (**z**) and (**dd**), for PBS (**ee-hh**) and PFF (**ii-ll**) injected rats, respectively. **mm-nn**) Quantification of C3 (**mm**) and pSyn (**nn**) fluorescence intensity in the ST. **oo**) Linear regression between pSyn and C3 fluorescence intensity in the ST of PFF injected rats. **pp-qq)** Quantification of percent area of ST occupied by C3+ (**pp**) and pSyn+ (**qq**) staining. **rr**) Linear regression between pSyn and C3 percent area staining in the ST of PFF injected rats. Data expressed as mean fold change (± standard deviation) from PBS controls (analyzed with unpaired t-test with Welch’s correction or simple linear regression analyses). Scale bars in (**h; dd)** are 250μm and apply to (**a-h; w-dd)**, respectively. Scale bars in (**p; ll)** are 50μm and apply to (**i-p; ee-ll)**, respectively.

However, while the upregulation of C3 was similar between the Cx and SNc (∼6 fold increase from controls in Cx versus ∼5 fold increase from controls in the SNc), there is significantly more pSyn pathology in the Cx versus the SNc (∼120 fold increase in the Cx versus ∼15 fold increase from controls in the SNc). Thus, there may be a disproportional upregulation of C3, relative to pSyn levels, in the SNc when compared to other regions. This is also true in the ST, however it is possible that some changes in the ST may be in response to the injectate itself (i.e. exogenous α-syn protein) as opposed to endogenous α-syn seeded into aggregates. As the SNc and Cx are physically separated from the injection site, this potential confound does not apply, suggesting an exaggerated complement response to Lewy body-like α-syn aggregates in the SNc. This differential response could potentially contribute to the selective vulnerability of nigral dopamine neurons to PD-associated pathology.

C3 activation can be stimulated by one of three complement activating pathways: classical, lectin and alternative (**Fig. 1a**). To determine which activating pathways are upregulated during the early phases of synucleinopathy, we quantified transcripts representing key targets within the different activation pathways using ddPCR. Specifically, we quantified genes representing the classical pathway (*C1qa*, *C1r* and *C4b*), the lectin pathway (mannan binding lectin serine protease 1 (*Masp1)* and mannose binding lectin 2 (*Mbl2)*) and the alternative pathway (complement factor D (*Cfd*) and complement factor B (*Cfb*)) in the ipsilateral SN of PBS and PFF injected rats (**Fig. 5a**). Expression of the classical pathway transcripts were all significantly increased in the SN of PFF injected rats compared to PBS controls. The expression of the lectin pathway target, *Masp1*, was significantly increased in the SN of PFF injected rats compared to PBS controls, however the magnitude of this increase (∼1.2-fold) was substantively lower than the magnitude of change in genes of the classical pathway (∼1.6-3.5 fold). Finally, expression of the alternative pathway targets were significantly increased (∼1.5 fold), albeit to a lesser extent than the classical pathway targets (**Fig. 5a**). These data suggest pathological α-syn upregulates genes associated with the classical pathway and, to a lesser extent, the alternative pathway.

**Fig. 5.**
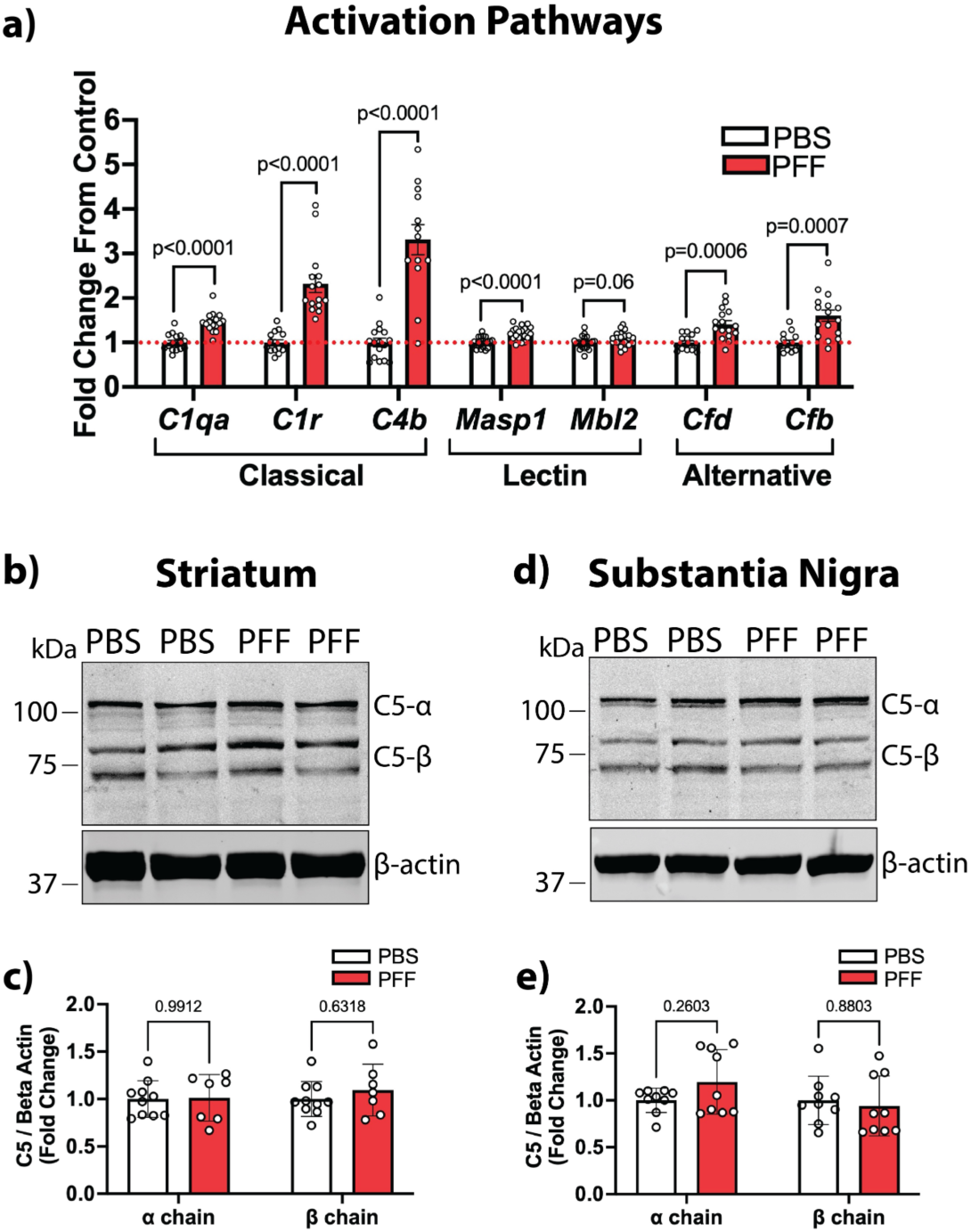
Preferential Upregulation of The Classical Pathway in the Substantia Nigra of a-syn PFF Injected Rats. Male and female rats (n=6-8/sex/group) received intra-striatal injec8ons of α-synuclein (α-syn) preformed fibrils (PFFs) or phosphate buffer saline (PBS) and were sacrificed 2-months post-injec8on. **a)** Droplet digital PCR (ddPCR) quantification of transcripts representing targets of the classical (*C1qa, C1r, C4b*), lectin (mannan-associated serine protease 1 (*Masp1),* mannose binding lectin 2 *(Mbl2*)) and alternative (complement factor D (*Cfd),* complement factor B *(Cfb*)) complement activation pathways in the ipsilateral substantia nigra (SN) from PBS and PFF injected rats. Data normalized to ribosomal protein L13 (*Rpl13*) and expressed as mean fold change (± standard deviation) from PBS controls (analyzed with multiple unpaired t-tests with Welch’s correction). **b-e)** Complement C5 (C5) immunoblotting analysis of the ipsilateral striatum (ST; **b-c**) and SN (**d-e**) from PBS and PFF injected rats (n=4-5/sex/group). **b)** Representative immunoblots of C5 and β-actin in the ST of PBS and PFF injected rats. **c)** Quantification of C5 α-chain (∼115 kDa) and α-chain (∼70, 76 kDa) normalized to β-actin in the ST. **d)** Representative immunoblot of C5 and β-actin in the SN of PBS and PFF injected rats. **e)** Quantification of C5 α-chain (∼115 kDa) and α-chain (∼70, 76 kDa) normalized to β-actin in the SN. Data are expressed as a mean fold change (± standard deviation) from the PBS control group (analyzed with multiple unpaired t tests comparing PBS versus PFF, with Holm Sidak multiple comparison test).

We next determined if there is activation or upregulation of the terminal pathway during the early phases of synucleinopathy in α-syn PFF injected rats. We used ddPCR to quantify the expression of the terminal pathway genes, *C5*, *C8a* and *C9* and immunoblotting to measure C5.

Transcripts encoding the terminal pathway genes, *C5*, *C8a* and *C9* were undetectable in the ST and SN of both PBS and PFF injected rats, yet these targets were detectable in the liver (**Supplemental Fig. 5 online resource**), confirming functionality of the ddPCR primer/probes pairs used. Despite the lack of quantifiable *C5* transcript in the brain, we detected C5 protein by immunoblot in both the ST and SN (**Fig. 5b-e**), but there was no difference in overall levels of C5 protein, or C5 activation products between PBS and PFF injected rats in either the ST or SN. These data indicate the terminal pathway is not activated during the early phases of synucleinopathy.

The complement system is regulated at multiple levels by complement regulatory proteins while specific receptors mediate some of many of the complement effector responses. We used ddPCR to quantify the expression of select genes representing the major complement regulatory proteins and complement receptors in the SN of PBS or α-syn PFF injected rats (**Fig. 6**). The specific complement regulatory genes quantified include the soluble complement regulators (clusterin (*Clu*) and complement factor H (*Cfh*)), the membrane bound complement regulators (complement receptor 1 like protein (*Cr1l*), *Cd55*, *Cd59* and *Cd46*), and the C1q binding proteins (neuronal pentraxin 1 (*Nptx1*), *Nptx2* and the NPTX-receptor (*Nptxr*)) [125].

**Fig. 6.**
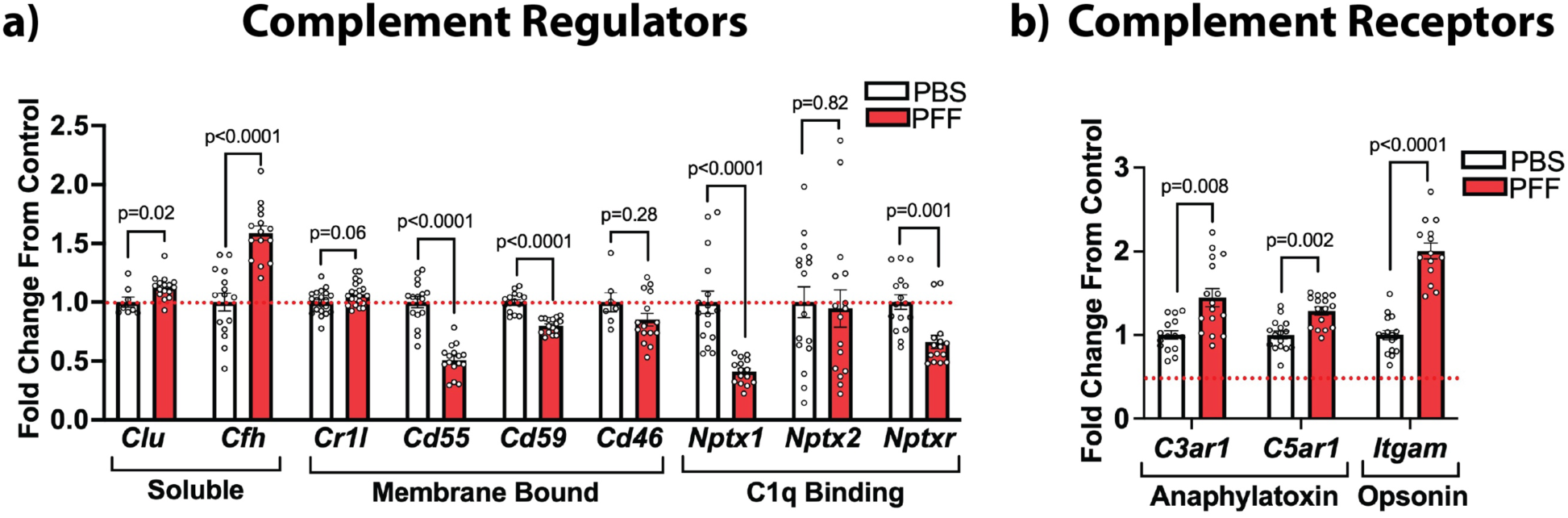
Complement Receptor and Regulator Expression are Altered in the Substantia Nigra of a-syn PFF Injected Rats. Male and female rats (n=6-8/sex/group) received intra-striatal injec8ons of α-synuclein (α-syn) preformed fibrils (PFFs) or phosphate buffer saline (PBS) and were sacrificed 2-months post-injec8on. **a)** Droplet digital PCR (ddPCR) quantification of transcripts encoding complement regulators, including the soluble regulators, clusterin (*Clu*) and complement factor H (*Cfh*); membrane bound regulators, complement receptor 1 like protein (*Cr1l*), *Cd55*, *Cd59* and *Cd46*; the C1q binding proteins, neuronal pentraxin 1 (*Nptx1*), neuronal pentraxin 2 (*Nptx2*), and the neuronal pentraxin receptor (*Nptxr*) in the ipsilateral SN of PBS and PFF injected rats. **b)** ddPCR quantification of transcripts encoding the anaphylatoxin receptors, *C3ar1*and *C5ar1*, and the *Itgam* gene encoding a subunit of the opsonin receptor, complement receptor 3 in the ipsilateral SN of PBS and PFF injected rats. Data normalized to ribosomal protein L13 (*Rpl13*) and expressed as a mean fold change (± standard deviation) from PBS controls (data analyzed with multiple unpaired t-tests and Holm Sidak multiple comparison test).

Expression of the soluble complement regulators, *Clu* and *Cfh*, were both significantly increased in the SN of PFF injected rats compared to PBS injected rats (**Fig. 6a**). In contrast, expression of the membrane bound complement regulators, *Cd55* and *Cd59*, and the C1q binding proteins, *Nptx1* and *Nptxr*, were significantly decreased in the SN of PFF injected rats (**Fig. 6a**). Next, we quantified the expression of genes encoding the anaphylatoxin receptors, *C3ar1* and *C5ar1*, as well as the *Itgam* gene, which encodes the integrin alpha M subunit of the opsonin receptor, complement receptor 3 (CR3). *C3aR1*, *C5aR1* and *Itgam* expression were all significantly increased in the SN of PFF injected rats compared to PBS injected controls (**Fig. 6b**). These data implicate the complement effector responses of opsonization (mediated through CR3) and inflammation (mediated through C3aR1 and C5aR1) and dysregulation of the complement system in response to synucleinopathy.

We next sought to assess changes in a representative complement receptor, for which we selected CR3, at the protein level using IF in midbrain tissue from PBS and α-syn PFF injected rats (**Fig. 7**). CR3 and IBA1 immunoreactivity were present throughout the ventral midbrain of PBS injected rats (**Fig. 7a-d, i-l**), however CR3 and IBA1 signal were dramatically increased in PFF injected rats, specifically in areas of the SNc containing pSyn aggregates, where CR3+ microglia were observed in direct apposition to pSyn+ aggregates (**Fig. 7e-h, m-p**), suggesting CR3+ microglial are targeting neurons containing pathological α-syn. Quantification of IBA1 (**Fig. 7q**) and CR3 (**Fig. 7s**) IF intensity and the percent area of the SNc occupied by IBA1+ (**Fig. 7t**) and CR3+ (**Fig. 7v**) staining were both significantly increased in the SNc of PFF injected rats, and this increase paralleled increased pSyn levels (**Fig. 7r, u**). These data indicate microglia upregulate expression of the opsonin receptor, CR3, in response to pathological α-syn.

**Fig. 7.**
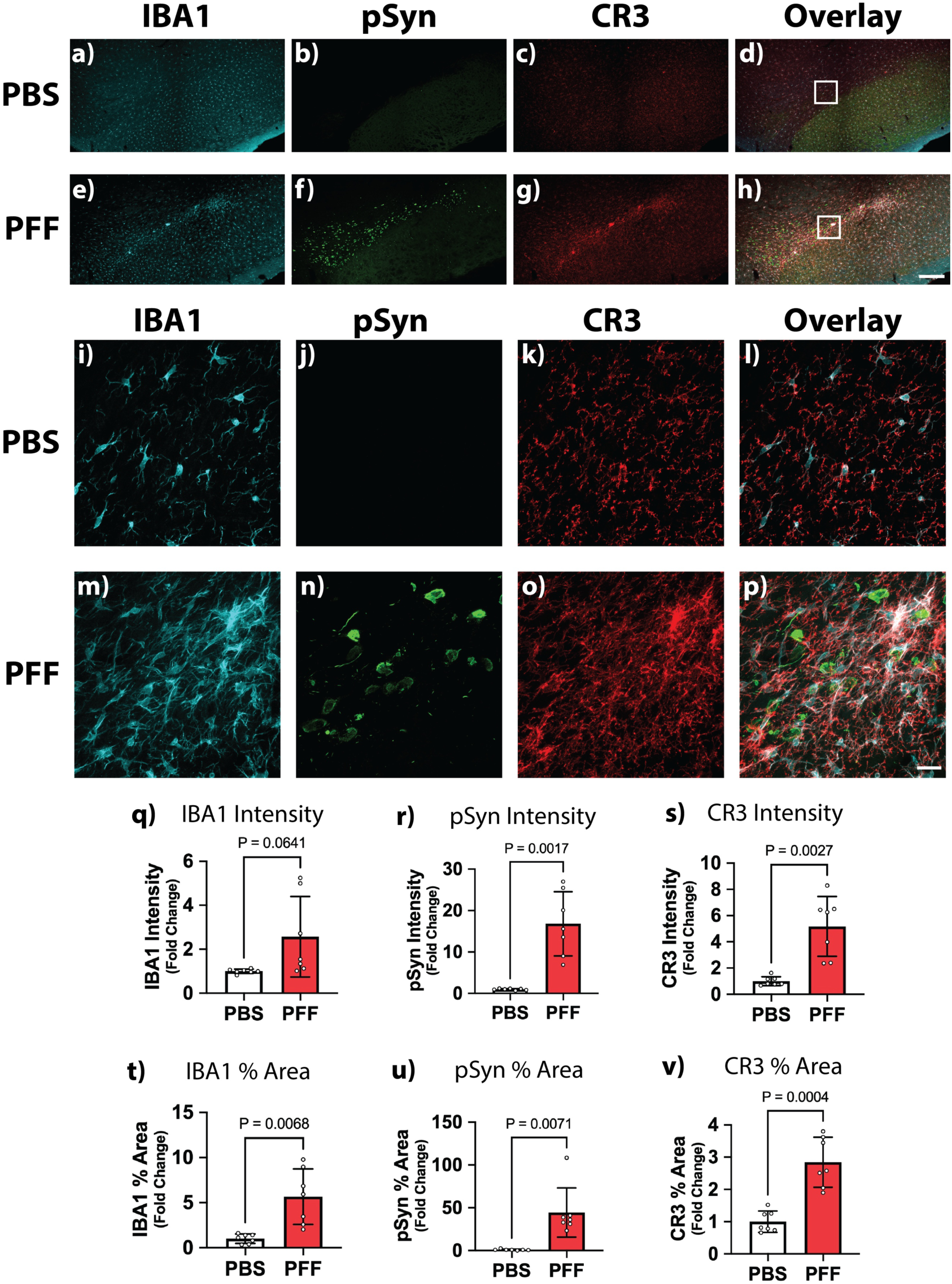
Microglial Complement Receptor 3 Expression is Increased in the Substantia Nigra of a-syn PFF Injected Rats. Male and female rats (n=6-7/group) received intra-striatal injec8ons of α-synuclein (α-syn) preformed fibrils (PFFs) or phosphate buffer saline (PBS) and were sacrificed 2-months post-injec8on. **a-h**) Representative low magnification immunofluorescent (IF) images of the microglial marker, ionized calcium binding adaptor molecule 1 (IBA; cyan), α-syn phosphorylated at serine 129 (pSyn; green) and complement receptor 3 (CR3; red) and the corresponding overlay image in the SNc of PBS (**a-d**) and PFF (**e-h**) injected rats. **i-p**) High magnification images corresponding to boxes in (**d**) and (**h**), for PBS (**i-l**) and PFF injected (**m-p**) rats, respectively. **q-s**) Quantification of IBA1 (**q**), pSyn (**r**) and CR3 (**s**) fluorescence intensity in the SNc. **t-v**) Quantification of the percent area of the SNc occupied by IBA1+ (**t**), pSyn+ (**u**) and CR3+ (**v**) staining. Data expressed as mean fold change (± standard deviation) from PBS controls (data analyzed with t-test with Welch’s correction). Scale bar in (**h)** is 250μm and applies to (**a-h**), scale bar in (**p)** is 50μm and applies to (**i-p**)

We also evaluated changes in a representative complement regulator, for which we selected Nptx1, at the protein level using IF in midbrain tissue from PBS and α-syn PFF injected rats (**Fig. 8**). Nptx1 immunoreactivity was decreased in the ventral midbrain of α-syn PFF injected rats (**Fig. 8e-h, m-p**) compared to PBS controls (**Fig. 8a-d, i-l**). High magnification imaging of Nptx1 in the SNc of PBS controls revealed diffuse Nptx1 immunoreactivity within the somata of nigral neurons combined with abundant Nptx1 puncta studding the neuropil throughout the SN (**Fig. 8i-l**). In contrast, Nptx1 immunoreactivity was visibly decreased in both nigral neuron somata and the SNc neuropil in α-syn PFF injected rats (**Fig. 8m-p**). Nigral neurons that were largely devoid of pSyn+ aggregates or contained smaller aggregates retained a strong, diffuse staining pattern that resembled Nptx1 immunoreactivity in nigral neurons of control animals (arrowheads in **Fig. 8m-p**). However, this diffuse staining pattern was almost entirely absent in neurons that contained pSyn+ aggregates, particularly aggregates that appeared more compact and mature where bright Nptx1 puncta colocalized with pSyn (arrows in **Fig. 8m-p, and Fig. 8q-s**). Quantification of Nptx1 fluorescence in the SNc confirmed a significant decrease in both Nptx1 IF intensity (**Fig. 8t**) and Nptx1 percent area staining (**Fig. 8v**) in α-syn PFF injected rats that was paralleled by significantly increased levels of pSyn in the SNc (**Fig. 8u, w**). These data suggest that, in addition to Nptx1 gene downregulation, Nptx1 protein may also be sequestered into α-syn aggregates during the development of synucleinopathy, further impairing its ability to regulate complement.

**Fig. 8.**
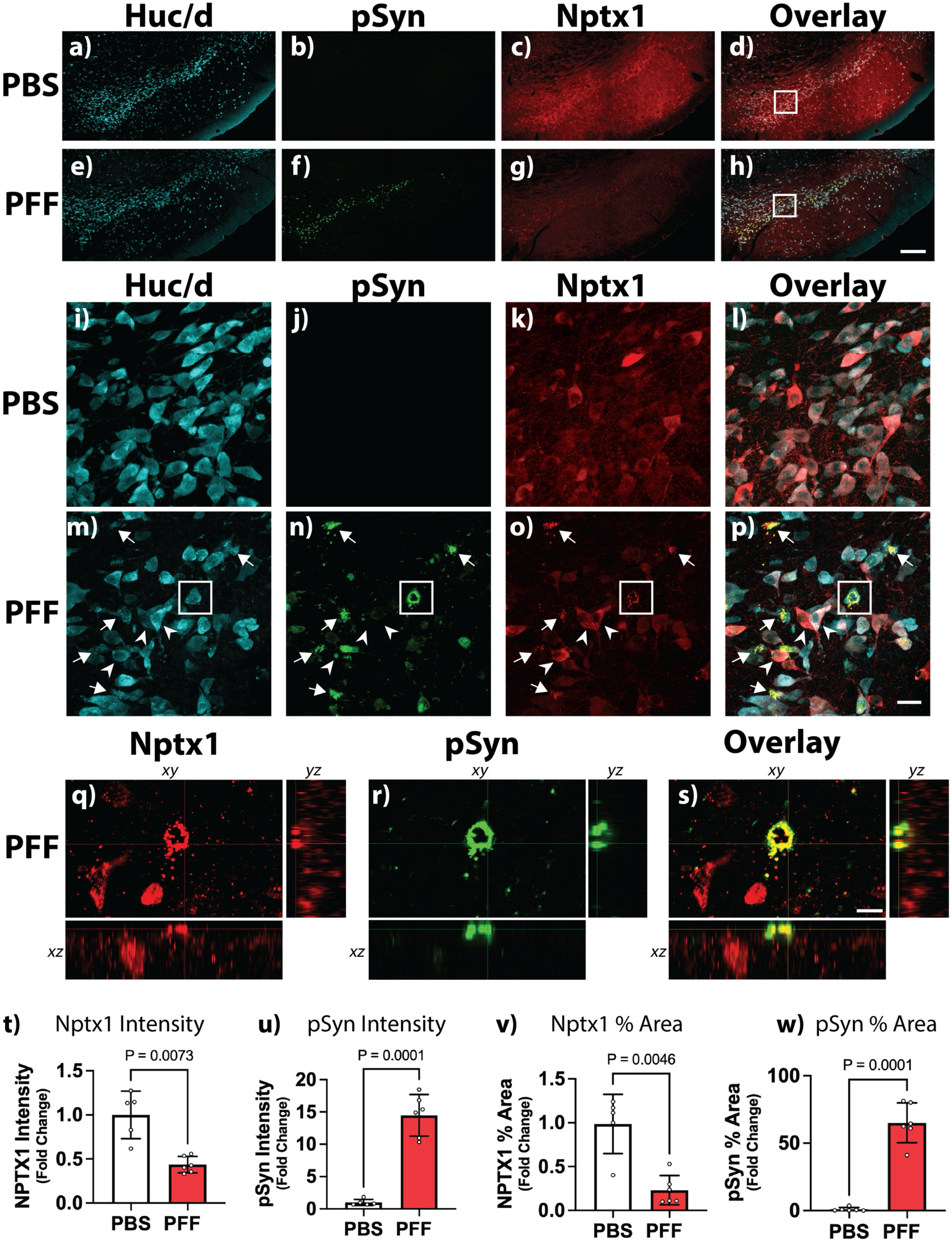
Neuronal Pentraxin 1 is Decreased and Localized to pSyn+ Inclusions in Nigral Neurons of a-syn PFF Injected Rats. Male and female rats (n=5-6/group) received intra-striatal injec8ons of α-synuclein (α-syn) preformed fibrils (PFFs) or phosphate buffer saline (PBS) and were sacrificed 2-months post-injec8on. **a-h**) Representative low magnification immunofluorescent (IF) images of the pan-neuronal marker Huc/d (cyan), α-syn phosphorylated at serine 129 (pSyn; green), neuronal pentraxin 1 (Nptx1; red) and the overlay image in the SNc of PBS (**a-d**) and PFF (**e-h**) injected rats. **i-p**) Representative high magnification images corresponding to boxes in (**d)** and (**h),** for PBS (**i-l**) and PFF injected (**m-p**) rats, respectively. Arrows in (**m-p)** indicate areas where Nptx1 co-localizes with pSyn+ aggregates. Arrowheads in (**m-p)** indicate neurons with diause Nptx1 immunoreactivity and minimal or absent pSyn+ inclusions. **Q-s**) Representative orthogonal view Z-stacks corresponding to the box in (**m-p)** showing colocalization between Nptx1 and pSyn in an aggregate. **t-u**) Quantification of Nptx1 (**t**) and pSyn (**u**) fluorescence intensity in the SNc. **v-w**) Quantification of the percent area of the SNc occupied by Nptx1+ (**v**) and pSyn+ (**w**) staining. Data expressed as mean fold change (± standard deviation) from PBS controls (data analyzed with t-test with Welch’s correction). Scale bar in (**h)** is 250μm and applies to (**a-h**), scale bar in (**p)** is 50μm and applies to (**i-p**), scale bar in (**s)** is 10μM and applies to (**q-s**)

Decreased complement regulatory proteins (CD55, CD59 and NPTX1) combined with the increased complement activation observed in the SN of PFF injected rats could render nigral dopamine neurons vulnerable to complement mediated attack. Complement activation is reported in the PD brain[63, 118], however prior reports did not specifically quantify complement regulatory proteins in PD affected brain regions. Thus, we quantified complement regulatory proteins in the PD SNc using postmortem midbrain tissue from neuropathologically confirmed control and PD brains (see Supplemental **Table 1 Online Resources** for neuropathologic and demographic data). We confirmed the presence of Lewy pathology in PD tissue using pSyn staining (**Fig. 9d-f**) and immunoblotted for the dopamine transporter (DAT) and tyrosine hydroxylase (TH) to confirm nigral neuron degeneration (**Fig. 9g-i**). We also quantified C1qa and C3/iC3b to determine the degree of complement upregulation/activation in the PD brain. Levels of C1qa were significantly increased in the SNc of PD brains (**Fig. 9j-k**). In contrast, there was no change in overall levels of C3 between the control and PD SNc (**Fig. 9l-m**). Quantification of iC3b by immunoblot trended (p=0.08) towards an increase while iC3b quantified by sELISA was significantly increased (**Fig. 9l-n**). Levels of CD55 (**Fig. 9o-p**) and NPTX1 (**Fig. 9w-x**) protein were significantly decreased in the SNc of PD brains compared to controls but no other significant changes were observed in complement regulators, though CD35 trended towards a decrease (p=0.08). These data, combined with the changes observed in the SNc of PFF injected animals, suggest that downregulation of CD55 and NPTX1 and increased levels of C1q and iC3b are central pathological changes associated with PD neuropathology.

**Fig. 9.**
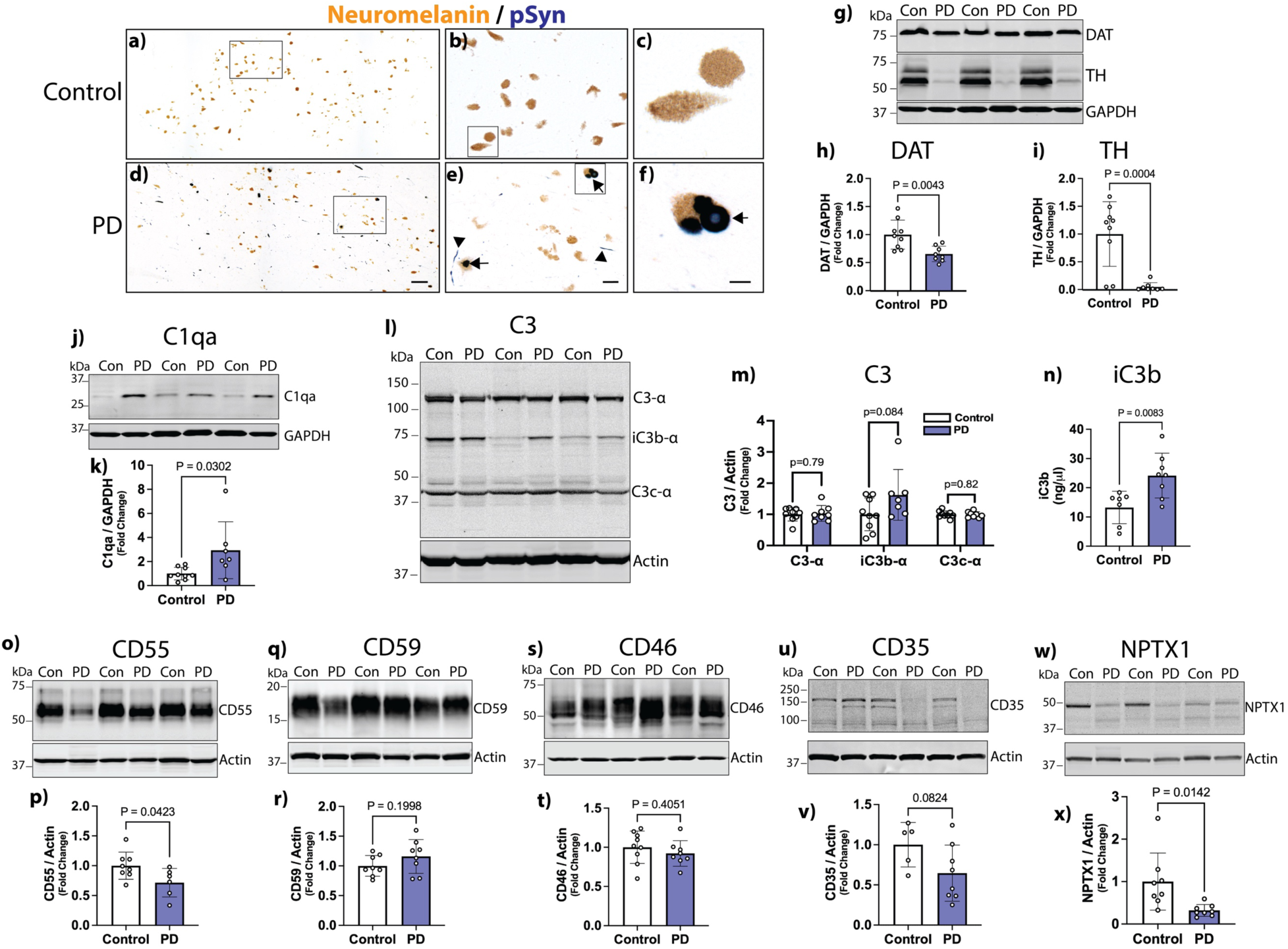
Expression of The Complement Regulatory Proteins, CD55 and NPTX1, are Decreased in the Parkinson’s Disease Substantia Nigra. Postmortem human midbrain 8ssue from age matched controls and neuropathologically confirmed PD pa8ents (n=5-8) was obtained from the Michigan Brain Bank. **a-f**) Representative images of immunostaining for α-synuclein (α-syn) phosphorylated at serine 129 (pSyn) in the SNc of control (**a-c**) and PD (**d-f**) tissue. **b, e**). High magnification images corresponding to the box in panels (**a, d**), respectively. **c, f**) High magnification images corresponding to the box in panels (**b, e),** respectively. Arrows in **(e-f)** indicate Lewy bodies while arrowheads in (**e-f)** indicate Lewy neurites. **g**) Representative immunoblot of the dopamine transporter (DAT), tyrosine hydroxylase (TH) and glyceraldehyde 3-phosphate dehydrogenase (GAPDH) in the SNc from control and PD tissue. **h-i**) Quantification of DAT (**h**) and TH (**i**) normalized to GAPDH. **J-k**) Representative immunoblot of C1qa and GAPDH (**J**) and quantification of C1qa normalized to GAPDH (**k**). **l-m**) Representative immunoblot of C3 and β-actin (**l**) and quantification of C3 α-chain (∼115 kDa), iC3b α-chain (∼67 kDa), and C3c α-chain (∼40 kDa) normalized to β-actin (**m**). Data in (**h, I, k, m**) are expressed as mean fold change (± standard deviation) from controls and were analyzed by t-test with Welch’s correction. **n**) Quantification of iC3b by sandwich ELISA. Data represent mean iC3b concentration (± standard deviation) analyzed by t-test with Welch’s correction. **o-x**) Representative immunoblots and quantification of CD55 (**o-p**), CD59 (**q-r**), CD46 (**s-t**), CD35 (**u-v**), neuronal pentraxin 1 (NPTX1; **panels w-x**) normalized to β-actin. Data expressed as mean fold change (± standard deviation) from controls and analyzed by t tests with Welch’s correction. Scale bar in (**d)** is 125μm and applies to (**a**), scale bar in (**e)** is 25μm and applies to (**b**), scale bar in (**f)** is 10μm and applies to (**c**).

Finally, we developed in vitro assays to test whether pathological α-syn directly binds C1q and activates complement (**Fig. 10**). C1q bound to human IgG (positive control) and human α-syn PFFs in a concentration dependent- and saturable manner. C1q also bound weakly to human α-syn monomers at higher concentrations but did not bind the negative control HSA (**Fig. 10a**). The average EC50 of C1q binding was 5.45nM for human IgG, 73.68nM for human α-syn PFFs, 167nM for human α-syn monomer and 153.8nM for HSA (**Fig. 10b**). The EC50 of IgG to binding to C1q was significantly lower than all other proteins, while the EC50 of α-syn PFF binding to C1q was significantly lower than HSA and α-syn monomers (**Fig. 10b**). There was no significant difference between the EC50 of α-syn monomer and the negative control, HSA, for C1q binding. Finally, to determine if the size of α-syn fibrils affects C1q binding, we performed the C1q binding assay with full length human α-syn PFFs and sonicated α-syn PFF, where both intact and sonicated human α-syn PFFs bound C1q in an identical manner (**Fig. 10c**).

**Fig. 10.**
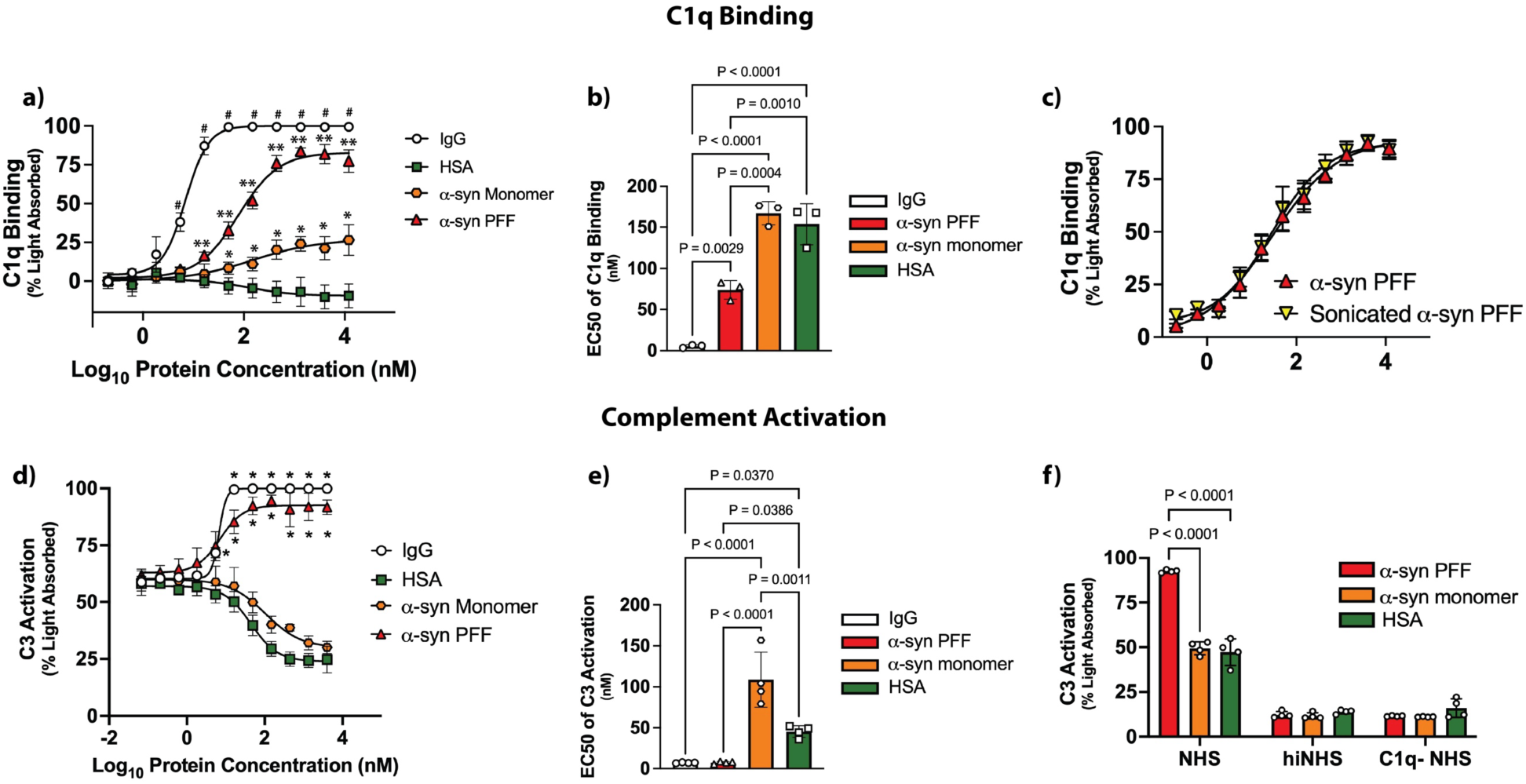
Aggregated a-synuclein Activates the Complement System in a C1q-Dependent Manner. **a**) C1q binding to human IgG (positive control), human serum albumin (HSA, negative control), human α-synuclein (α-syn) monomers and human α-syn preformed fibrils (PFFs), expressed as mean percent light absorbed (± standard deviation, n=3 biological replicates) analyzed by two-way ANOVA and Tukey’s multiple comparison. ^#^ significantly diaerent from HSA, α-syn monomer and PFFs within same concentration; ** significantly diaerent from HSA and α-syn monomer within same concentration; * significantly diaerent from HSA within same concentration. **b**) C1q binding EC_50_ for human IgG, α-syn monomer, α-syn PFFs, and HAS, as mean EC_50_ values (± standard deviation, n=3 biological replicates) analyzed with a one-Way ANOVA and Tukey’s multiple comparison. **c**) C1q binding to intact α-syn PFFs or sonicated α-syn PFFs, expressed as mean percent light absorbed (± standard deviation, n=3 biological replicates) analyzed with a two-way ANOVA and Tukey’s multiple comparison. **d**) Complement activation (indexed by formation of C3b/iC3b) by human IgG (positive control), HSA (negative control), α-syn monomer and α-syn PFFs, expressed as mean percent light absorbed (± standard deviation, n=4 biological replicates) analyzed by two-way ANOVA and Tukey’s multiple comparison. * Significantly diaerent from HSA and α-syn within same concentration. **e**) EC_50_ for complement activation by human IgG, α-syn monomer, α-syn PFFs, and HAS, expressed as mean EC_50_ values (± standard deviation, 4 independent biological replicates) analyzed with a one-Way ANOVA and Tukey’s multiple comparison. **f**) Complement activation assay using α-syn monomer, α-syn PFFs or HSA using normal human serum (NHS), heat inactivated NHS (hiNHS), or C1q depleted NHS (C1q-NHS), expressed as means percent light abosrbed (± standard deviation, n=4 biological replicates) analyzed by one-way ANOVA and Tukey’s multiple comparison.

We next sought to determine if aggregated α-syn activates the complement system in a cell free system. Human IgG (positive control) and human α-syn PFFs increased the formation of C3b/iC3b (cleaved/activated forms of C3) in a concentration dependent manner that was saturable. In contrast, human α-syn monomer and HSA did not increase C3b/iC3b levels (**Fig. 10d**), where levels of activated C3 decreased at the higher concentrations of HSA and α-syn monomer compared to blank wells containing only blocking buffer (2% bovine serum albumin). The average EC50 for C3 activation was 6.9 nM for human IgG, 7.2 nM for human α-syn PFFs, 108.5 nM for human α-syn monomer, and 45.2 for HSA (**Fig. 10e**). The EC50 for IgG and human α-syn PFF induced C3 activation was significantly lower than human α-syn monomer and HSA, while the EC50 for C3 activation in the α-syn monomer group was significantly higher than all other groups (**Fig. 10e**). Finally, to determine if complement activation by pathological α-syn is mediated through the classical activation pathway we repeated the complement activation assay using C1q depleted normal human serum (C1q-NHS). Depletion of C1q from serum completely prevented C3 activation in response to α-syn PFFs, down to the level of the negative control, heat inactivated NHS (hiNHS; **Fig. 10f**). Together, these data show that pathological α-syn can directly bind C1q and activate the complement system in a C1q-dependent.

## Discussion

The goal of the current study was to determine if pathological α-syn activates the complement system prior to neurodegeneration. Uncontrolled complement activation is increasingly recognized as a potential driver of neurodegeneration in PD, and numerous studies report complement upregulation and activation in the PD brain and α-syn-based preclinical models of PD[12, 19, 40, 42, 46, 62, 64, 66, 77, 81, 100, 108, 118, 119, 126]. However, all previous studies have investigated complement activation at a time when significant neurodegeneration has already occurred. As such, whether complement activation is triggered directly by pathological α-syn has remained unclear.

Given the abundant evidence linking complement activation to the degenerative process in PD, we hypothesized that, if complement contributes to toxicity, significant complement activation and upregulation should occur long before death of nigral neurons. To test this hypothesis, we quantified complement gene expression and complement activation during the early phases of synucleinopathy in the rat α-syn PFF model. We report; 1) robust upregulation and activation of C3 that correlates with levels of pSyn across several brain regions, 2) a preferential upregulation of classical pathway components, 3) increased expression of soluble complement regulators, accompanied by decreased expression of select membrane bound- and C1q-binding complement regulators, 4) a significant upregulation of anaphylatoxin-and opsonin receptors, and 5) microglia as the primary cellular source of C3 and CR3 in the rat brain during the early phases of synucleinopathy.

Importantly, all the above findings occurred in the SNc during peak α-syn aggregate formation, but months prior to nigral neuron degeneration, supporting the conclusion that synucleinopathy induces the activation of complement, independent of cell death. We further validated key findings in postmortem human PD SNc tissue and found a significantly decreased CD55 and NPTX1 protein combined with increased levels of C1q and iC3b protein. Finally, we demonstrate that aggregated, but not monomeric, α-syn directly activates the complement system in a C1q-dependent manner. Taken together, these findings reveal complement activation during the early phases of synucleinopathy and provide compelling evidence that α-syn-induced complement dysregulation may contribute to degeneration in PD and related synucleinopathies.

### Pathological α-syn Directly Activates Complement

One of the most salient conclusions from the current study is that aggregated forms of α-syn can directly activate the complement system. Indirect evidence supporting this conclusion include increased expression of numerous complement genes and elevated levels of activated C3 in the SNc of α-syn PFF injected rats, findings consistent with previous reports of complement upregulation and/or activation in α-syn based preclinical models of PD[12, 64, 77, 100, 119, 126]. However, the current study and our recent report [100] uniquely demonstrate complement upregulation and activation during the early phases of synucleinopathy, indicating these changes arise in response to aggregated α-syn, independent of degeneration. Consistent with this interpretation, both C3 expression and levels of activated C3 are increased in the SNc of α-syn PFF injected rats, where C3 upregulation significantly correlates with pSyn burden.

Microglia expressing C3 and CR3 are similarly increased in the SNc of α-syn PFF injected rats, where they localize to pSyn+ inclusions and correlate with pSyn levels. Finally, C3+ microglia also spatially overlap with pSyn+ aggregates in the Cx and ST and correlate with levels of pSyn in the Cx. The striking spatial correspondence between pSyn+ aggregates and C3/CR3+ microglia, combined with the significant correlation between levels of C3 and pSyn load, strongly implicate complement upregulation in response to aggregated α-syn. Importantly, the brain regions where C3 significantly correlated to pSyn burden (SNc and Cx) are anatomically distant from the injection site, indicating complement upregulation and activation in these regions are driven by aggregates composed of endogenous misfolded α-syn, rather than injected recombinant protein or injection related injury.

These data are corroborated by direct evidence demonstrating complement activation in response to aggregated α-syn in our *in vitro* complement assays. In these assays, human α-syn aggregates bound purified human C1q and induced a concentration-dependent deposition of activated C3. Because these experiments were conducted in a completely cell free system, they provide evidence that aggregated α-syn is sufficient to activate the complement system. Notably, complement activation was specific to aggregated α-syn, as human α-syn monomer did not stimulate complement activation. Finally, the complement response to aggregated α-syn was C1q dependent.

C1q is a soluble pattern recognition receptor (PRR) that initiates the classical complement pathway. As a PRR, C1q recognizes common molecular motifs associated with cellular damage, termed “damage associated molecular patterns” (DAMPs), or motifs associated with pathogens, termed “pathogen associated molecular patterns” (PAMPs). C1q can also recognize motifs associated with neurodegenerative disease pathology, such as β-amyloid [44], which are referred to as “neurodegeneration associated molecular patterns” (NAMPs). With this view, aggregated α-syn is likely recognized as a NAMP or DAMP by C1q, thereby stimulating complement activation. Supporting this, aggregated α-syn also binds numerous other PRRs expressed on innate immune cells (reviewed in [31]).

Aggregated α-syn represents a rare, pathological protein conformation associated with cellular dysfunction and toxicity. As such, C1q and the complement system may target aggregated α-syn for phagocytic clearance to prevent or limit toxicity. In contrast, monomeric α-syn is one of the most abundant proteins in the brain and performs many essential cellular functions [6, 41]. Moreover, monomeric α-syn and complement components, including C1q, coexist in many body tissues, including blood and brain[5, 41]. Accordingly, if soluble, monomeric α-syn could activate complement, one would expect a basal level of complement activation in healthy individuals. However, this is not the case, as chronic complement activation is typically associated with disease and/or inflammatory or immunological disorders, not normal physiology.

The results from our *in vitro* complement activation assays are consistent with some previous findings but not others[28, 50]. For example, [50] found that full length, monomeric α-syn did not activate the complement system, whereas the alternatively spliced isoform, α-syn 112, did. In contrast, [28] found that full length α-syn, but not the truncated α-syn 1-95, induced complement activation. Our results align closely with those of [50], however the reason for these discrepancies are unclear. One possibility is that some complement activation attributed to monomeric α-syn in previous studies were driven by *de novo* aggregate formation.

α-syn is an intrinsically disordered protein that can spontaneously misfold into oligomers and fibrils at RT. In our experiments, monomer samples are thawed on ice and centrifuged to prevent aggregate formation and remove contaminating aggregates [82]. Without these precautions, we have observed spontaneous formation of *de novo* α-syn aggregates capable of eliciting an immune response (unpublished data). It is possible that without these precautions there may have been contaminating aggregates that caused complement activation in previous studies. This hypothesis is in line with the increased complement activation reported with α-syn 112, an isoform with accelerated aggregation kinetics [65, 84]. However, it would not account for the lack of complement activation reported with α-syn 1-95, as C-terminal truncation of α-syn also accelerates aggregation [95]. Alternatively, methodological differences, such as the complement activation product quantified (C3b/iC3b versus C4b versus C5b-9), may explain these discrepant findings. Future studies that rigorously control for *de novo* aggregate formation are needed to definitively determine whether monomeric α-syn can activate the complement system.

### Synucleinopathy Primarily Activates Complement Through the Classical Pathway

Data from the current study indicate pathological α-syn stimulates complement activation through the classical pathway, leading to increased opsonization and anaphylatoxin signaling. We found significant upregulation of targets from both the classical and alternative pathways. However, the magnitude of classical pathway gene upregulation was greatest.

Expression of the lectin pathway protease, *Masp1*, was also statistically increased, though the magnitude of change was minimal. Together, these findings indicate synucleinopathy drives a broad elevation in complement gene expression, with a pronounced bias toward classical pathway activation. Consistent with this conclusion, purified human C1q bound aggregated human α-syn and depletion of C1q from human serum completely abolished complement activation in response to pathological α-syn. However, the upregulation of alternative pathway components, such as Factor D and Factor B, suggests the alternative pathway may amplify the complement response to synucleinopathy once initiated through C1q.

The pattern of classical pathway upregulation and activation observed in α-syn PFF injected rats aligns with complement activation in the PD brain. Numerous studies utilizing a variety of methods (including gene expression, transcriptomics, proteomics, and histology) document upregulation and activation of classical pathway targets in postmortem PD tissue[19, 42, 62, 66, 118]. Among these altered complement targets, C4 and C1-complex proteins are the most consistently elevated and/or activated, mirroring findings from the present study.

Additionally, levels of activated C4 are increased in the brains of patients with Dementia with Lewy Body[40, 108], indicating complement activation is not unique to PD but a more general response to synucleinopathy. Finally, our results align with prior work demonstrating that C1q is required for α-syn-induced complement activation[28]. These data indicate complement activation is a common response to aggregated α-syn and support the conclusion that C1q recognizes pathological α-syn as a DAMP or NAMP, thereby triggering complement activation through the classical pathway.

### Early Synucleinopathy Stimulates Phagocytosis and Anaphylatoxin Signaling

Following activation, current data indicates that opsonization and anaphylatoxin signaling are the primary complement effector responses enacted during the early phases of synucleinopathy. Regardless of the initial activation pathway stimulated, complement activation ultimately results in the cleavage of C3 into a small peptide named C3a and a larger fragment named C3b. C3a is a potent anaphylatoxin that increases local inflammation by recruiting and activating immune cells. C3b is an opsonin that covalently binds targets to mark them for phagocytic clearance by cells expressing CR1. However, C3b is rapidly cleaved by Factor I and Factor H to generate iC3b, which is also a functional opsonin that recruits phagocytes through interaction with CR3 and CR4.

In the current study, levels of the opsonin, iC3b, were significantly increased in both the ST and SNc of α-syn PFF injected rats. Additionally, expression of the *Itgam* gene, which encodes a subunit of CR3, was one of the most highly upregulated receptor genes in the SNc of α-syn PFF injected rats. Microglia expressing CR3 were increased in the SNc of PFF injected rats and, like C3+ microglia, localized to the precise location of pSyn+ aggregates. Interestingly, in addition to binding iC3b, CR3 is also a PRR that can directly bind α-syn, thereby facilitating microglial uptake, and genetic deletion of CR3 prevents microglial activation in response to pathological α-syn [37, 45, 111, 123]. These data strongly implicate CR3 and complement mediated phagocytosis in response to synucleinopathy.

Currently it is unclear if α-syn aggregates or neurons containing these aggregates are labeled with opsonins and/or targeted by CR3 expressing phagocytes in α-syn PFF injected rats. We did not observe colocalization of C3 with pSyn+ aggregates at the time point investigated.

However, the antibody used to detect C3 in the current study was not specific to activated forms of C3. Thus, it is possible that α-syn aggregates or the neurons containing aggregates were labeled with C3b/iC3b but were not detected because the reagents used were not specifically sensitive to activated C3. Alternatively, it is possible that complement opsonins may label α-syn aggregates or nigral neurons at later timepoints in the degenerative cascade.

Supporting this conclusion, both neuromelanin neurons and Lewy bodies label with iC3b in postmortem PD tissue [62]. Additional experiments are needed to determine if α-syn aggregates are opsonized during the early phases of synucleinopathy. However, we did observe C3+ microglia and CR3+ microglia in direct apposition to neurons containing pSyn+ aggregates. These complement expressing microglia appeared to be extending processes around neurons containing α-syn aggregates, suggesting they may be targeting synucleinopathy-affected neurons for phagocytic clearance. Taken together, these results support the hypothesis that activated complement targets aggregated α-syn for phagocytic clearance.

The current study also implicates anaphylatoxin signaling in the complement response to synucleinopathy. The anaphylatoxins, C3a and C5a, are potent inflammatory signaling molecules that function through interactions with their cognate receptors, C3aR and C5aR. C3aR and C5aR are expressed on multiple CNS cell types, most prominently microglia, but also astrocytes, endothelial cells, perivascular macrophages, and under specific conditions, neurons. Activation of these receptors induce a wide range of proinflammatory responses, including glial priming and activation, chemotaxis, cytokine release, vascular changes and increased phagocytosis, while activation of neuronally expressed anaphylatoxin receptors is directly toxic to cultured neurons, including mesencephalic dopamine neurons [34, 80, 112]. C3aR and C5aR expression were significantly increased in the SNc of α-syn PFF injected rats compared to controls, indicating increased C3a/C5a signaling during the early phases of synucleinopathy. Although C3aR upregulation has been reported in the SN of A53T α-syn transgenic mice[12], our findings extend this observation by demonstrating anaphylatoxin receptor upregulation in a second, mechanistically distinct model of α-syn pathology. Apart from the A53T study, increased C3aR/C5aR expression has not been widely documented in PD, making this a novel aspect of the PD complement response.

Anaphylatoxin signaling, particularly through C5aR, triggers downstream expression of immune genes that shift glial towards a reactive, pro-inflammatory phenotype[27, 33, 61, 88]. We recently identified a distinct suite of immune genes upregulated in reactive microglia and astrocytes in the SN during the early phases of synucleinopathy[77, 100]. Interestingly, many of these genes or their associated networks overlap with anaphylatoxin-responsive transcriptional programs described in the context of AD[27, 61, 88], raising the possibility that C3a/C5a signaling regulates the conversion of homeostatic glia to the reactive state observed in PD. This convergence suggests anaphylatoxin receptor signaling may represent a shared inflammatory pathway across neurodegenerative diseases and highlights their potential as therapeutic targets in synucleinopathies. Future studies testing the ability of C3aR/C5aR inhibition to prevent inflammation and toxicity in models of PD are warranted.

### Complement Dysregulation During Early Synucleinopathy

Finally, results from the current study implicate complement regulator dysfunction as a potential source of neurotoxicity in synucleinopathies. Complement regulators are a diverse group of proteins that restrain complement activation at different levels of the cascade to protect self-cells from excessive inflammation, opsonization and membrane attack.

Complement regulators fall broadly into three categories, soluble fluid-phase regulators, membrane bound regulators, and C1q binding proteins. In the current study we observed a consistent pattern of dysregulation characterized by upregulation of fluid phase regulators, paralleled by a downregulation of specific membrane bound regulators and C1q-binding proteins. This imbalance suggests that key homeostatic “checkpoints” may be compromised in early synucleinopathy, potentially lowering the threshold for complement-mediated glial activation and/or neuronal injury.

Fluid phase regulators are secreted proteins that circulate in plasma, CSF and extracellular space. These include Serping1 (inhibits C1r and C1s), Complement Factor H (catalyzes breakdown of C3b), and clusterin, (binds components of the terminal pathway). In the current study, expression of Factor H and clusterin were significantly increased in the SN of α-syn PFF injected rats, indicating a non-cell autonomous upregulation of fluid phase regulators during early synucleinopathy. This finding is consistent with our recent report of increased Serping 1 expression in the SN during the aggregation phase of the PFF model [100]. As α-syn directly activated the classical pathway in a C1q-dependent manner, increasing opsonin deposition and phagocytic receptor expression, increased Serping 1 and Factor H likely represent compensatory responses aimed at limiting mistargeted phagocytosis during heightened complement activation.

Clusterin upregulation may serve a dual role in the response to synucleinopathy. In its canonical complement regulatory role, clusterin binds soluble C7, C8β and C9 to inhibit MAC formation, however, clusterin also acts as a molecular chaperone to prevent aggregation and promote clearance of misfolded proteins [10]. Thus, clusterin upregulation in α-syn PFF injected rats may be an attempt to mitigate α-syn aggregation. Numerous lines of evidence support this [86]idea. For example, clusterin directly binds aggregated α-syn to reduce cellular uptake and aggregation in cultured cells overexpressing α-syn [24, 57], while clusterin co-localizes to lewy bodies in DLB and glial cytoplasmic inclusions in MSA[86]. Clusterin is increased in nigral neurons of AAV α-syn and α-syn PFF treated mice[24], closely mirroring results from the current study. Clusterin levels are increased in the CSF and blood of PD patients (reviewed in[10] and clusterin upregulation is highest in early-stage PD patients [110]. Finally, polymorphisms in the gene encoding clusterin significantly increase the risk of PD[59], indicating loss of clusterin function may be detrimental. These data indicate clusterin upregulation is a conserved, early-stage protective response to synucleinopathy.

Membrane bound complement regulators are cell surface proteins that protect host cells from unintended complement mediated damage by locally inhibiting different steps of the complement cascade. These regulators include CD46 (Co-factor to inactivate C3b and C4b), CD55 (catalyzes dissociation of C3 and C5 convertase), CD59 (prevents incorporation of C9 into the C5b-8 complex), CD35 (accelerates dissociation of C3 and C5 convertases and cofactor to inactivate C3b and C4b). In the current study, CD55 and CD59 expression were significantly decreased in the SN of α-syn PFF injected rats during early synucleinopathy, and CD55 and NPTX1 were significantly decreased in the PD SNc. These results align with prior reports of CD55 downregulation in PD and related models. CD55 protein is decreased in the ST of α-syn PFF injected mice[119], while meta-analyses report decreased CD55 expression in the PD SNc [46].

Additionally, a recent study found increased CD55 transcripts in peripheral blood but decreased CD55 protein in the CSF of PD patients, where CD55 levels correlate with motor and cognitive impairment and help distinguish PD from controls [8]. Finally, few studies have linked CD59 to PD. One study found increased CD59 protein in the PD SNc affected by both Lewy pathology and neuronal loss compared with SNc affected by neuronal loss alone [81]. Collectively, these data support CD55 dysfunction as a central event in PD pathobiology and the current study extends previous findings by demonstrating CD55 and CD59 downregulation during the early stage of synucleinopathy.

Lastly, the current results also implicate dysregulation of C1q-binding complement regulators in the response to synucleinopathy. Specifically, expression of *Nptx1* and the *Nptxr* were significantly decreased in the SN of α-syn PFF injected rats during the early phases of synucleinopathy. Although originally characterized for their role in synapse organization and plasticity, the neuronal pentraxins have more recently emerged as important modulators of complement in the CNS through their ability to bind C1q and inhibit classical pathway activation [52, 125]. In doing so, the neuronal pentraxins play a major role in regulating complement mediated synaptic pruning. Decreased nigral expression of Nptx1 during synucleinopathy is a novel finding, however previous reports link the neuronal pentraxins to PD. NPTX2 is upregulated in the PD SNc where it colocalizes with α-syn in Lewy bodies and neurites [67], mirroring the colocalization of NPTX1 in α-syn aggregates observed in the SNc of PFF injected rats in the current study. NPTX1 is increased in the PD hippocampus [114], and all neuronal pentraxins are decreased in the CSF of patients with PD, MSA and progressive supranuclear palsy[70]. Importantly, CSF neuronal pentraxin levels correlate with cognitive impairment, dopaminergic degeneration and motor impairment in PD, implicating the neuronal pentraxins in disease pathophysiology and as biomarkers[70]. Collectively, these findings and the current results implicate neuronal pentraxin dysregulation in the response to synucleinopathy.

The neuronal pentraxins regulate complement mediated synaptic pruning, and loss of this function is implicated in the synaptic degeneration associated with tauopathy[125]. Accordingly, decreased nigrostriatal NPTX1 during synucleinopathy could decrease complement inhibition at the synapse, leading to increased C1q deposition, opsonization, and microglial engulfment. Loss of synaptic complement regulation provides a plausible mechanistic link between pathological α-syn accumulation, aberrant complement activation, and early synaptic dysfunction in PD. Notably, nigrostriatal degeneration occurs in a “dying-back” fashion in PD, where degeneration of striatal terminals precedes degeneration of nigral somata[9, 36, 51, 105, 117]. Accordingly, loss of neuronal pentraxin function at striatal terminals, combined with increased complement activation in the PD brain[15, 19, 40, 42, 62, 66, 108, 118], could enable unregulated complement synaptic pruning of striatal terminals.

More broadly, the current data implicate dysfunction or downregulation of multiple classes of complement regulators in the response to PD pathology. These findings support a model in which complement dysregulation, driven both by increased activation and impaired regulation, contributes to synaptic vulnerability and neurodegeneration in synucleinopathies. Accordingly, therapeutic strategies aimed at maintaining or restoring complement regulator function could provide neuroprotection in PD and other synucleinopathies.

### Limitations of the Current Study

There are several limitations to the current study that should be considered when interpreting results. First, we were unable to comprehensively profile all components of the complement system. The complement system comprises over 50 proteins, and due to the targeted nature of the investigation, it was not feasible to profile every component. Accordingly, we may have missed important changes in the complement response to early synucleinopathy. In line with this issue, we could not localize the cell source of all differentially expressed complement components in α-syn PFF injected rats. Determining the cellular source of these changes could be critical for understanding how complement dysregulation contributes to neurodegeneration in synucleinopathies and future studies should be performed to address this issue. A major technical barrier we encountered was a lack of reliable reagents capable of detecting rat complement proteins with sufficient specificity and sensitivity. For example, we detected increased levels of iC3b (an activated fragment of C3) in α-syn PFF injected rats, however we were unable to reliably quantify activated fragments of most other complement proteins due to the lack of cleavage specific antibodies, or assays lacking the sensitivity necessary to detect activated complement in rat brain. As a result, we inferred complement activation or the initiation of different complement effector responses based on coordinated changes in complement gene expression rather than on the detection of specific activation products. Despite these limitations, multiple independent lines of evidence, including increased iC3b, robust upregulation of anaphylatoxin receptors, altered expression of complement regulators, strong spatial correlations with α-syn pathology, and cell-free assays demonstrating C1q-dependent activation by aggregated α-syn converge to support the conclusion that complement activation occurs during the early phases of synucleinopathy.

Lastly, the current study used of late-stage postmortem PD tissue to validate molecular changes identified during the early phases of synucleinopathy in α-syn PFF–injected rats. As such, the molecular changes observed in postmortem tissue could reflect late-stage consequences of neurodegeneration, rather than the primary response to synucleinopathy. This temporal disconnect prevents a direct comparison to the early complement changes observed in preclinical models. However, comparing changes between α-syn PFF injected rats and postmortem PD tissue yields important insights into the time course of complement dysregulation. For example, we detected robust upregulation of C3 and increased levels of iC3b in α-syn PFF injected rats. In contrast, overall levels of C3 were not changed in the PD SNc, while levels of iC3b were increased when quantified by sELISA. These discrepancies support the notion that C3 upregulation is an early response to synucleinopathy, which likely diminishes after significant neurodegeneration occurs. In contrast, CD55 and NPTX1 were significantly downregulated during early synucleinopathy in α-syn PFF injected rats and in late-stage postmortem PD tissue, indicating that CD55 and NPTX1 downregulation in response to synucleinopathy is conserved across species and disease stages. Collectively, these comparative findings suggest that while some of the complement response to synucleinopathy is transient and stage dependent, others, such as CD55/NPTX1 downregulation, may be durable changes that contribute to neurodegeneration.

### Conclusions

We conclude that synucleinopathy stimulates complement activation and dysregulation prior to overt degeneration. Complement activation during early synucleinopathy in α-syn PFF injected rats includes upregulation of both the classical and alternative pathways, and upregulation and activation of complement C3 in the absence of terminal pathway upregulation. We determined that microglia are the primary cellular source of complement C3 in the rat brain during early synucleinopathy and microglia upregulate C3 across multiple brain regions affected by synucleinopathy. Increased opsonization, phagocytosis and anaphylatoxin signaling are the primary complement effector responses initiated during early synucleinopathy. Further, early synucleinopathy causes downregulation of CD55, CD59 and Nptx1 in the SNc of α-syn PFF injected rats, and CD55 and NPTX1 are also decreased in the PD SNc, indicating that complement dysregulation is a durable change that persists through the pathological time course. Finally, we conclude that aggregated α-syn, but not monomeric α-syn, directly activates the classical complement system in a C1q-dependent fashion. The robust complement activation and regulator dysfunction observed during the early phases of synucleinopathy reported herein could render neurons susceptible to complement mediated attack. Taken together, the current results suggest that restoring normal complement regulation could be a viable therapeutic strategy for PD and other synucleinopathies.

## List of Abbreviations

Alpha-synuclein (α-syn), Association for Assessment and Accreditation of Laboratory Animal Care (AAALAC), Alternative pathway C3 convertase (C3bBb), BL21(DE3) *E. coli* expression strain (DE3), Bovine serum albumin (BSA), Carboxypeptidase-N (CPN), C1 inhibitor (C1-INH), Classical/lectin pathway C3 convertase (C4b2a), Clusterin (CLU/Clu), Complement component 3 (C3), Complement component 5 (C5), Complement factor B (Cfb), Complement factor D (Cfd), Complement factor H (Cfh), Complement receptor 1-like protein (Cr1l), Complement receptor 3 (CR3), C4 binding protein (C4BP), Dementia with Lewy bodies (DLB), Dopamine transporter (DAT), Droplet digital PCR (ddPCR), Dulbecco’s phosphate-buffered saline (dPBS), Electron microscopy (EM), Ethylenediaminetetraacetic acid (EDTA), Factor H (FH), Factor I (FI), Glyceraldehyde-3-phosphate dehydrogenase (GAPDH), Heat-inactivated normal human serum (hiNHS), Institutional Animal Care and Use Committee (IACUC), Isopropyl β-D-1-thiogalactopyranoside (IPTG), Mannose-binding lectin (MBL), Mannose-binding lectin 2 (Mbl2), Mannan-binding lectin serine protease 1 (Masp1), Membrane attack complex (MAC), Multiple system atrophy (MSA), Neuronal pentraxin 1 (Nptx1), Neuronal pentraxin 2 (Nptx2), Neuronal pentraxin receptor (Nptxr), Neuronal pentraxins (NPTXs), Normal human serum (NHS), Parkinson’s disease (PD), Phosphate-buffered saline (PBS), Phosphate-buffered saline with Tween-20 (PBS-T), Phosphorylated alpha-synuclein (pSyn), Pre-formed fibrils (PFFs), Ribosomal protein L13 (Rpl13), Sandwich enzyme-linked immunosorbent assay (sELISA), Sodium dodecyl sulfate (SDS), Striatum (ST), Substantia nigra (SN), Substantia nigra pars compacta (SNc), Substantia nigra pars reticulata (SNr), Thioflavin-T (ThT), Tris-buffered saline (TBS), Tris-buffered saline with Triton X-100 (TBS-Tx), Tris-buffered saline with Tween-20 (TBS-T), Tyrosine hydroxylase (TH), Vitronectin (VTN), Wild type (WT).

## Declarations

### Ethics Approval

All postmortem human tissue used in the current study was de-identified. At no point was any of the tissue identified or linked to any identifying records of any kind. As such, this tissue is determined Not Human Subjects under the Michigan State University Study ID STUDY00009841. All procedures involving animals were performed in accordance with federal, state and institutional guidelines approved by the Michigan State University Institutional Animal Care and Use Committee (IACUC) under protocol number PROTO202400182.

### Consent for Publication

Not Applicable.

### Availability of Data and Materials

The datasets generated and analyzed in the current study will be made available upon reasonable request to the corresponding author.

### Authors Contributions

Conception of the study: MJB; Design of the study: MJB; Production of α-syn PFFs: JRP, KCL; Acquisition of study results: MJB, HK, AK, SP, MG, AB, ACS, CES, CJK, JRP, KSC, NCK; Interpretation of study results: MJB; Drafting and revisions of manuscript: HK, MJB, NMK, CES, CJK. All authors read and approved of the final manuscript.

## Acknowledgements

The authors would like to thank the Michigan Brain Bank and the Michigan Alzheimer’s Disease Research Center for providing the postmortem human tissue, and the people who generously donated the tissue used in the current study.

