## Supplemental Figures for "Complement Dysregulation During the Early Phases of Synucleinopathy"

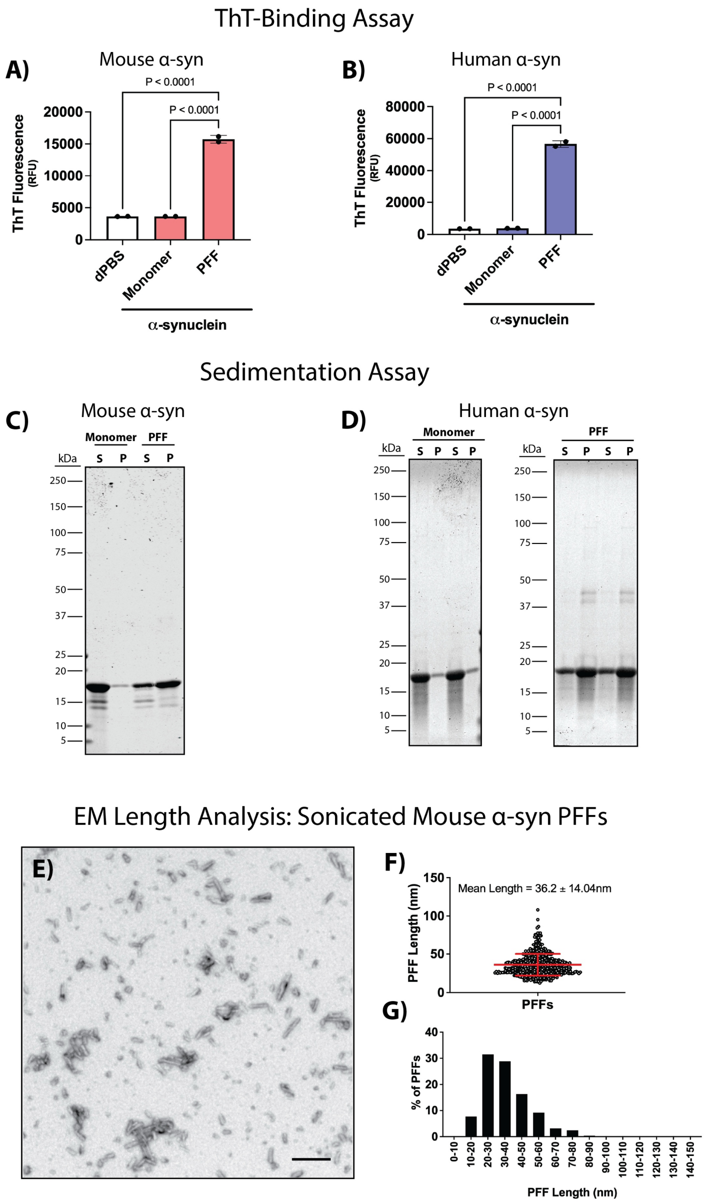


**Supplemental Fig. 1: α-syn Preformed Fibril Quality Control.** Quality control was performed on recombinant, full length mouse and human α-synuclein (α-syn) pre-formed fibrils (PFFs). **A-B**) β-sheet amyloid structures in assembled recombinant full length α-syn PFFs, α-syn monomers and Dulbecco's phosphate buffered saline (dPBS) vehicle were quantified with Thioflavin T (ThT). Recombinant mouse α-syn PFFs (**A**) and human PFFs (**B**) bound significantly more ThT than α-syn monomers and dPBS vehicle control (n=2 replicates, data analyzed by one way ANOVA with Tukey's multiple comparison test). **C-D**) A sedimentation assay was performed to detect high molecular weight α-syn species. Mouse (**C**) and human (**D**) α-syn monomers resolved primarily in the soluble fraction (S) while the majority of α-syn PFF protein resolved in the pellet fraction (P). **E-F)** Quantification of sonicated mouse α-syn PFF length by electron microscopy (EM). **E**) Representative electron micrograph showing negatively stained, sonicated mouse α-syn PFFs used for stereotactic injection. **F**) Size (nm) distribution of sonicated α-syn PFFs quantified from EM images. **G**) Size distribution (nm) of sonicated mouse α-syn PFFs expressed as a percentage of total PFFs that fall into 10nm bins. The mean length of sonicated moues α-syn PFFs was 36 ± 14nm and > 93% of all fibrils measured were ≤ 50nm. Scale bar in panel (**E**) is 200nm.

**A)**

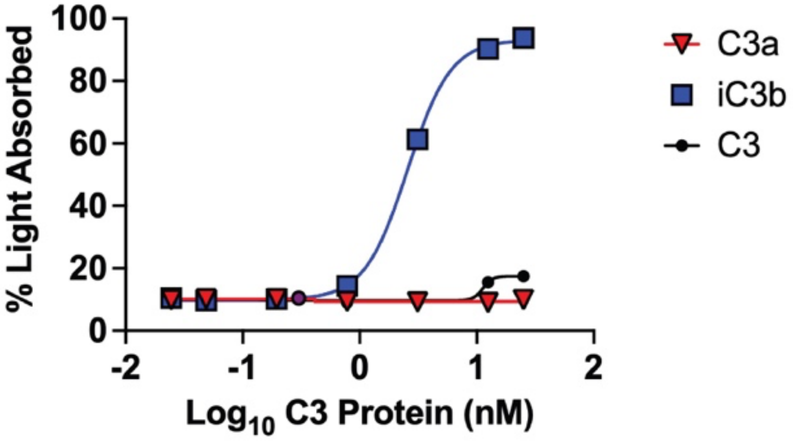


**B)**

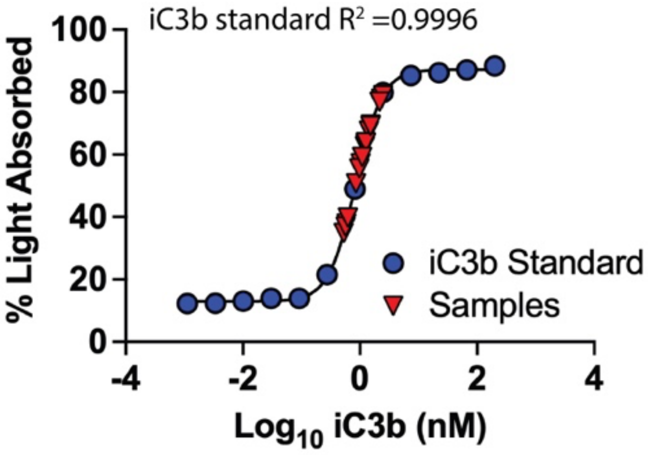


**Supplemental Fig 2: Human iC3b Sandwich ELISA. A**) The specificity of the assay for human iC3b was validated using purified human C3 (black lines with circles), human iC3b (blue lines with squares) and human C3a (red lines with triangles) ranging from 50nM to 0.048nM. **B**) Plot of iC3b standards (blue circles) and human brain lysate samples (red triangles) that were used to quantify iC3b in the substantia nigra of control and PD brains (these samples and standards correspond to **Figure 9N**). Note that the human samples are within the linear range of the standards.

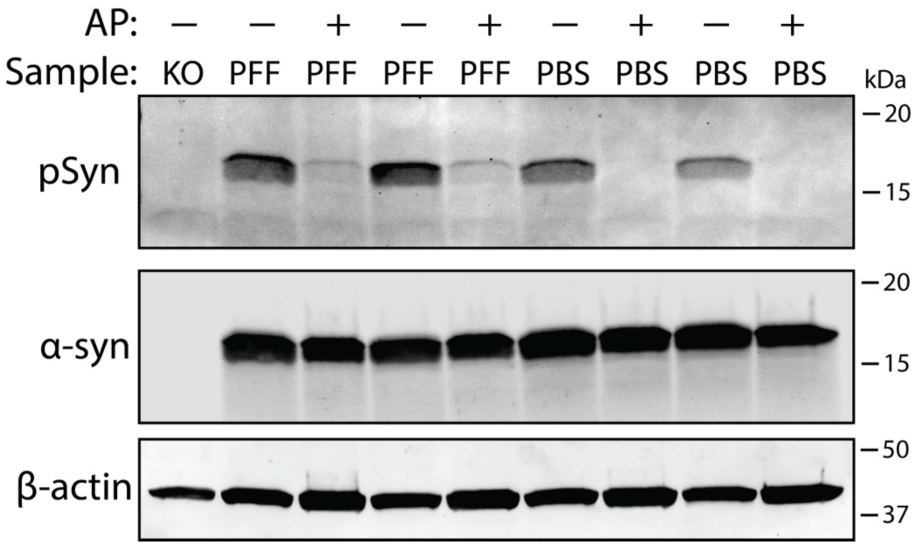


**Supplemental Fig 3. Testing the Specificity of the Antibody Used to Biochemically Quantify pSyn.** Striatal lysates from $\alpha$-synuclein ($\alpha$-syn) pre-formed fibril (PFF) or phosphate buffered saline (PBS) injected rats (n=4-5/group; 2-months post-injection), and untreated $\alpha$-syn germline knockout mice (KO; 6 months of age, n=3) were incubated with (+) or without (-) alkaline phosphatase (AP) to dephosphorylate $\alpha$-syn. Blots were probed with a pan $\alpha$-synuclein antibody ($\alpha$-syn) or a phospho-Serine 129 specific $\alpha$-syn (pSyn) antibody. Beta-actin was used as a loading control. Neither antibody detected protein bands in $\alpha$-syn KO lysates. The pSyn antibody detected a protein band resolving at ~17kDa, which was increased in the ST of $\alpha$-syn PFF injected rats compared to PBS controls. De-phosphorylation of lysates eliminated pSyn antibody signal in ST lysate of PBS treated rats and robustly decreased detection in PFF animals.

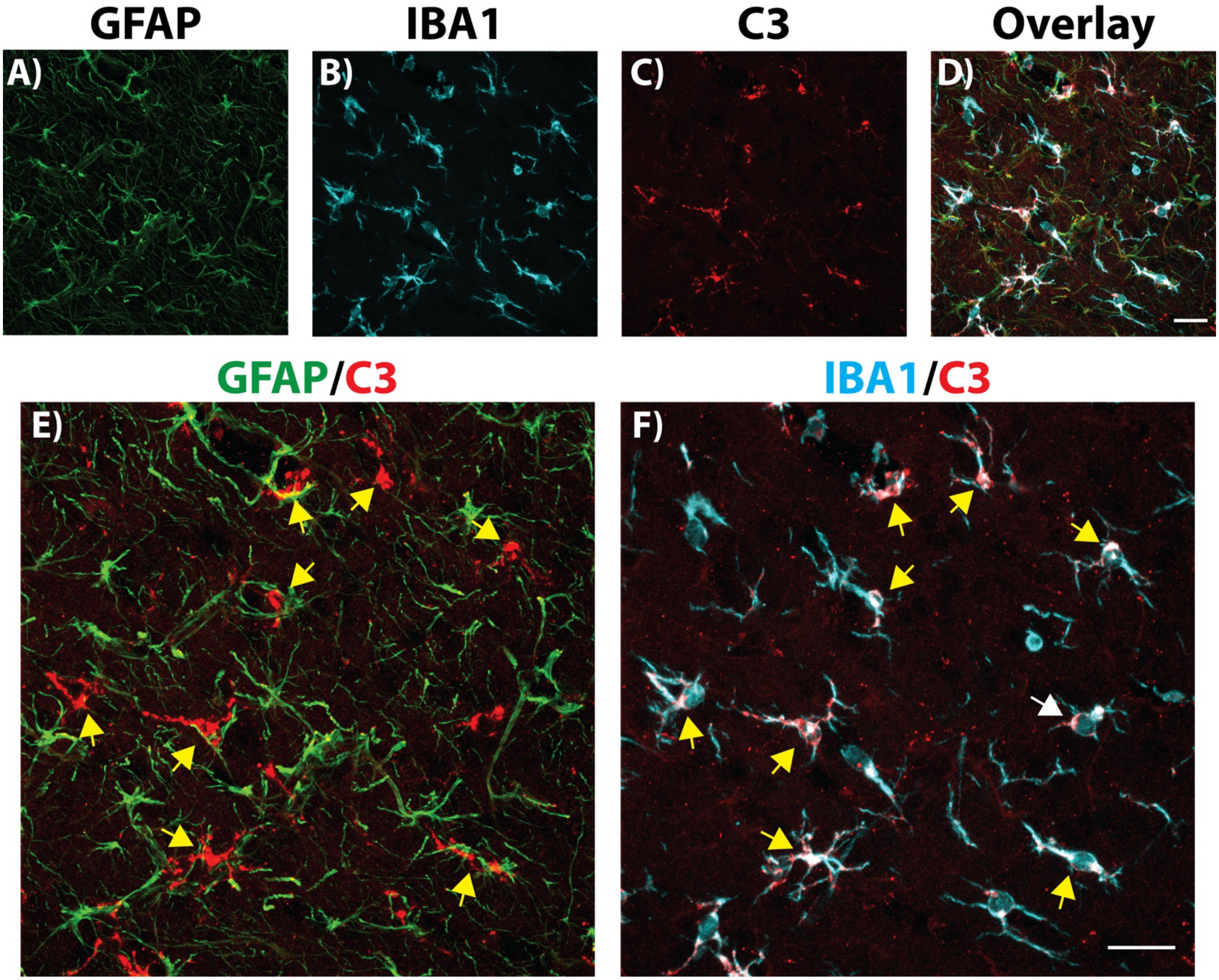


**Supplemental Fig. 4: Microglia are the Primary Cellular Source of Complement C3 in the Rat Brain.** Tissue from the ipsilateral cortex of α-synuclein (α-syn) pre-formed fibril (PFF) injected rats was processed for immunofluorescent (IF) detection of astrocytes (**panel A**; glial fibrillary acidic protein, GFAP; green), microglia (**panel B**; ionized calcium binding adaptor molecule 1, IBA1; cyan) and complement component 3 (**panel C**; C3; red). Enlarged overlay images of GFAP and C3, or IBA1 and C3 are shown in panels (**E**) and (**F**), respectively. Arrows in (**F**) show colocalization of C3 and IBA1 signal. Scale bars in (**D)** and (**F)** are 25μm and apply to (**A-C)** and **E** , respectively.

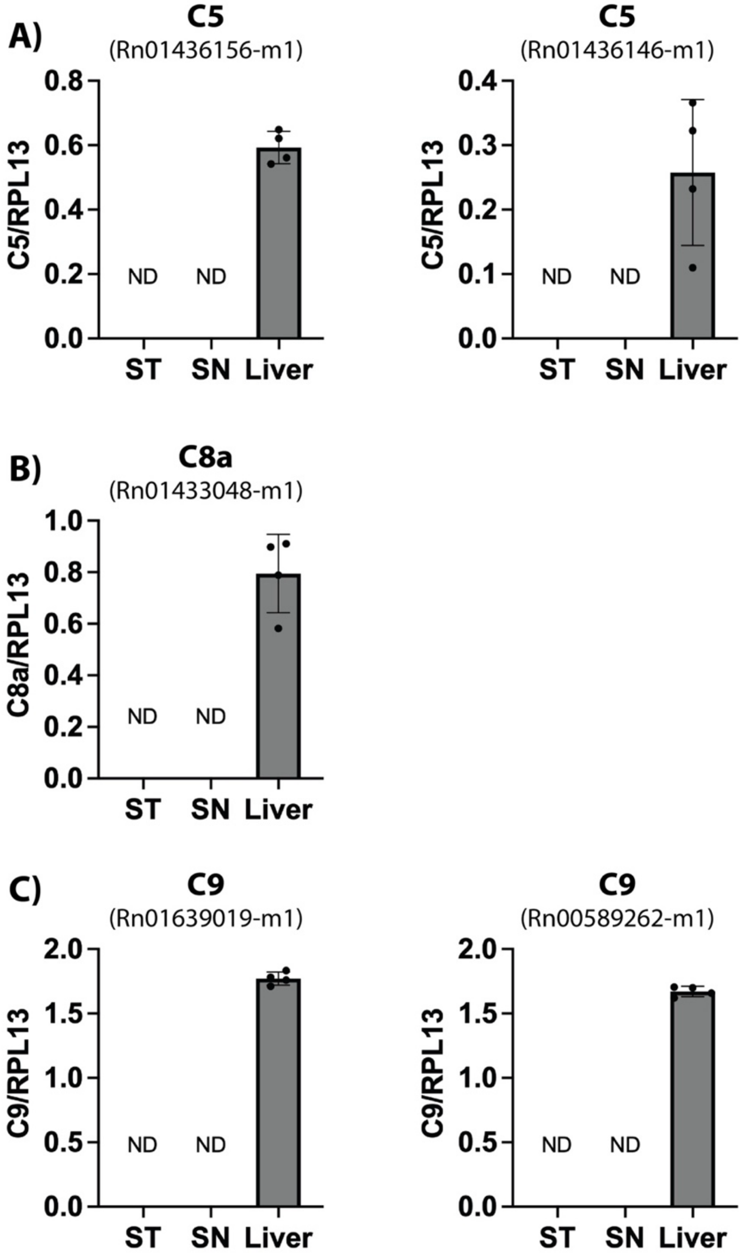


**Supplemental Fig. 5: Expression of Genes Encoding the Terminal Complement Pathway Targets, C5, C8a and C9, in the Striatum and Substantia Nigra of $\alpha$-syn PFF Injected Rats.** Male and female rats received intra-striatal injections of $\alpha$-synuclein ($\alpha$-syn) pre-formed fibrils (PFFs) (n=4-5/group) and were euthanized 2-months post-injection **A-C**)The liver, ipsilateral substantia nigra (SN) and striatum (ST) from $\alpha$-syn PFF injected animals were analyzed for transcripts representing different targets in the terminal complement pathway using droplet digital PCR. Quantification of complement *C5* (**A**), *C8a* (**B**), and *C9* (**C**) in the ST, SN and liver of PFF injected rats. Data represent the mean target gene levels ($\pm$ standard deviation) normalized to a housekeeping gene, ribosomal protein L13 (Rpl13). Catalogue numbers of the specific primer/probes sets are noted above each respective histogram. Not detected (ND).

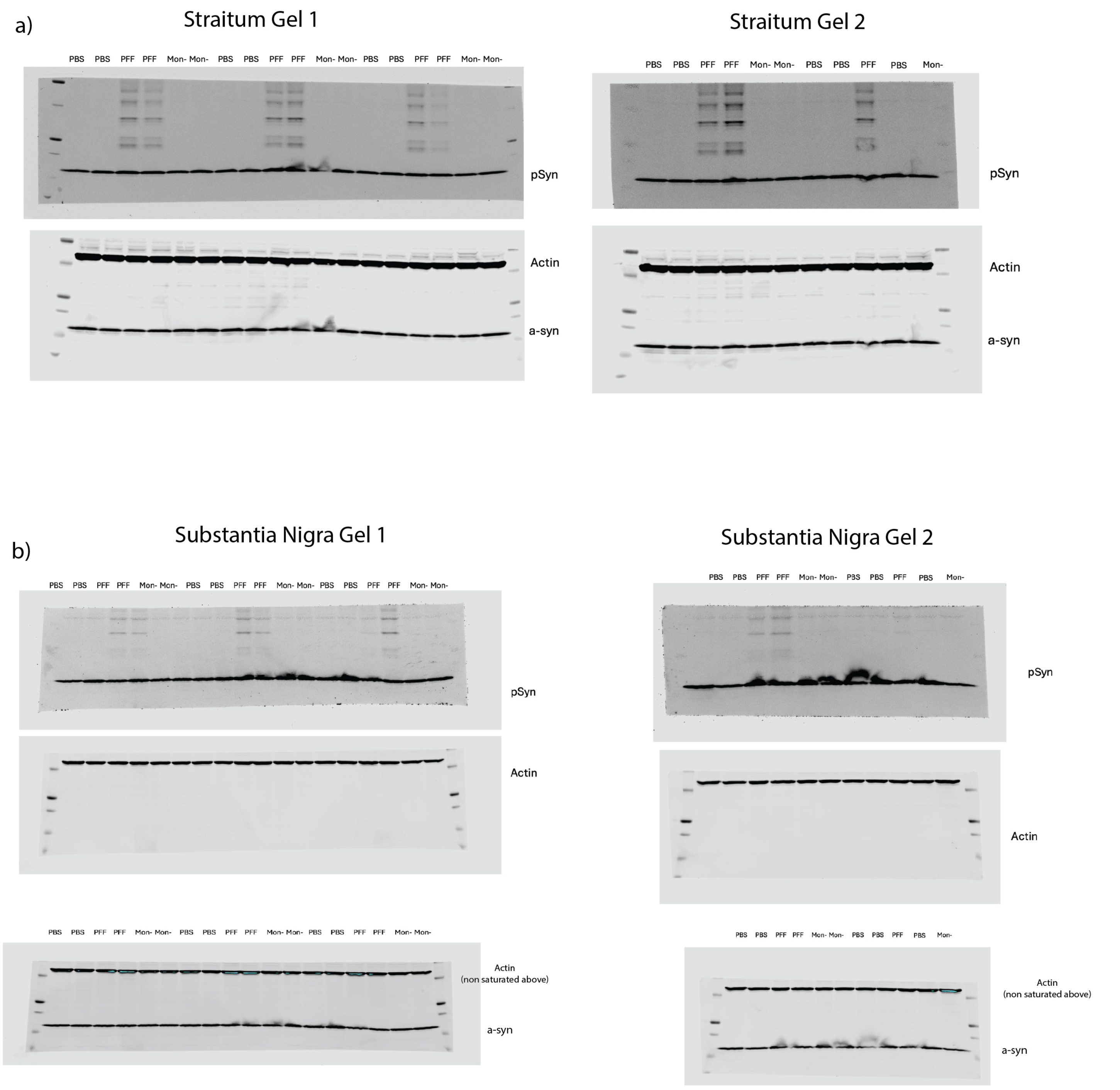


**Supplemental Fig. 6**: Uncropped Western blots corresponding to Figure 2 in main manuscript, showing serine 129 phosphorylated $\alpha$-synuclein (pSyn), $\alpha$-synuclein ($\alpha$-syn) and $\beta$-actin loading control in the Striatum (a) and substantia nigra (b) of PBS and $\alpha$-syn PFF injected rats.

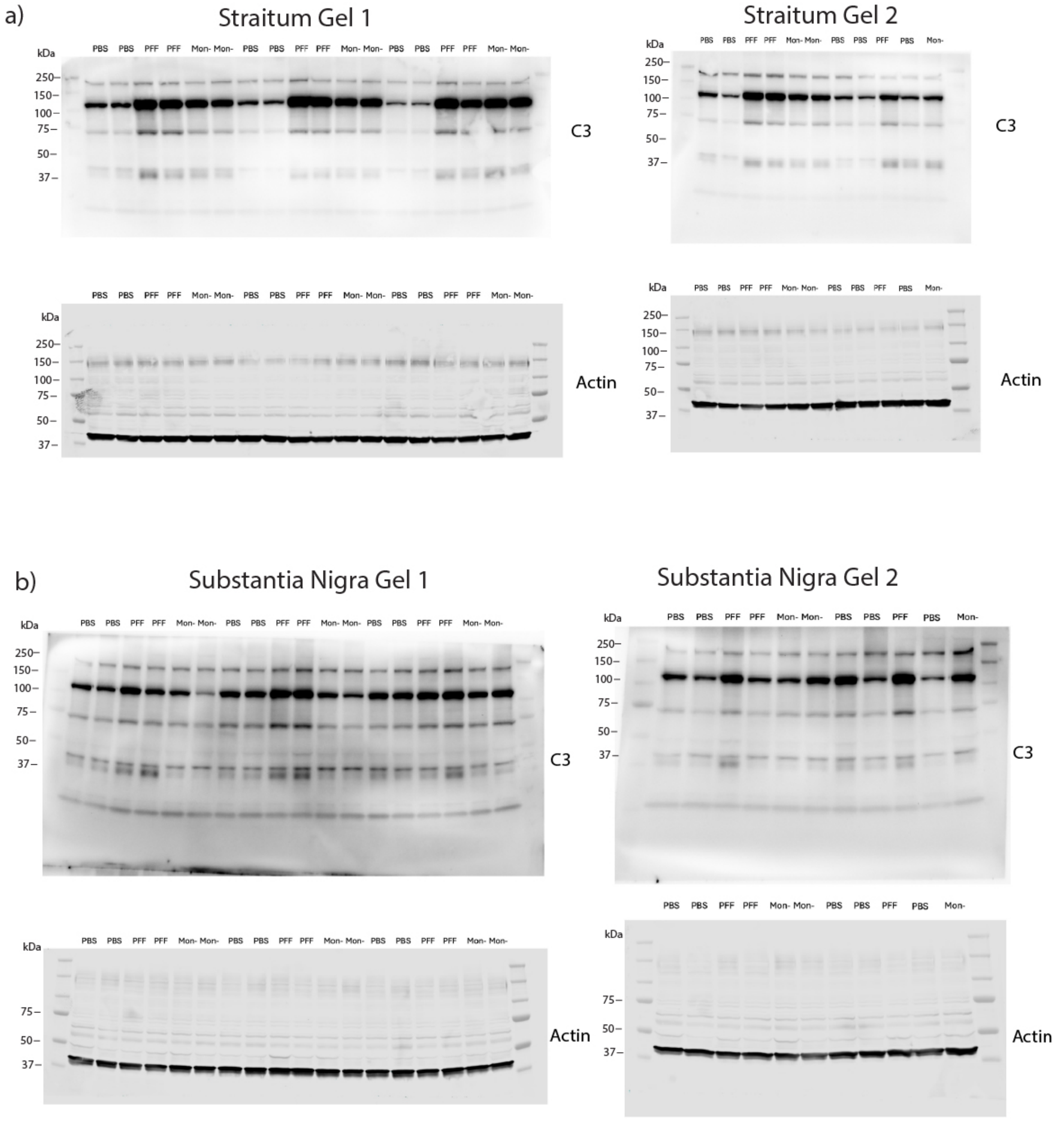


**Supplemental Fig. 7**: Uncropped Western blots corresponding to Figure 2 in main manuscript, showing complement C3 and β-actin loading control in the Striatum (a) and substantia nigra (b) of PBS and α-syn PFF injected rats.

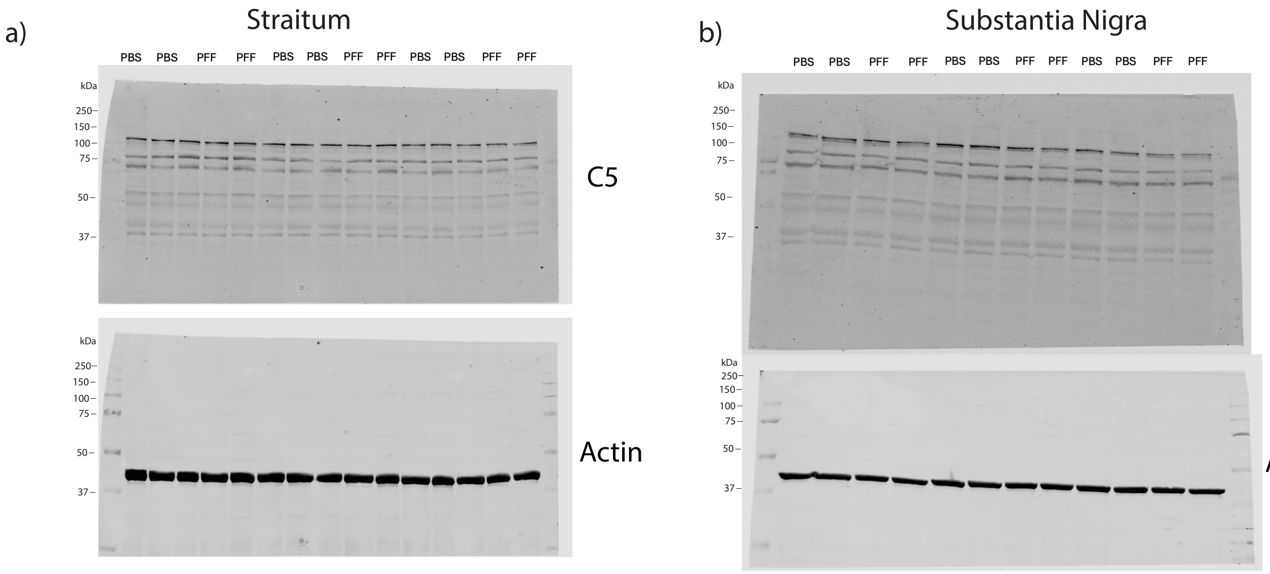


**Supplemental Fig. 8**: Uncropped Western blots corresponding to Figure 5 in main manuscript, showing complement C5 and β-actin loading control in the striatum (a) and substantia nigra (b) of PBS and α-syn PFF injected rats.

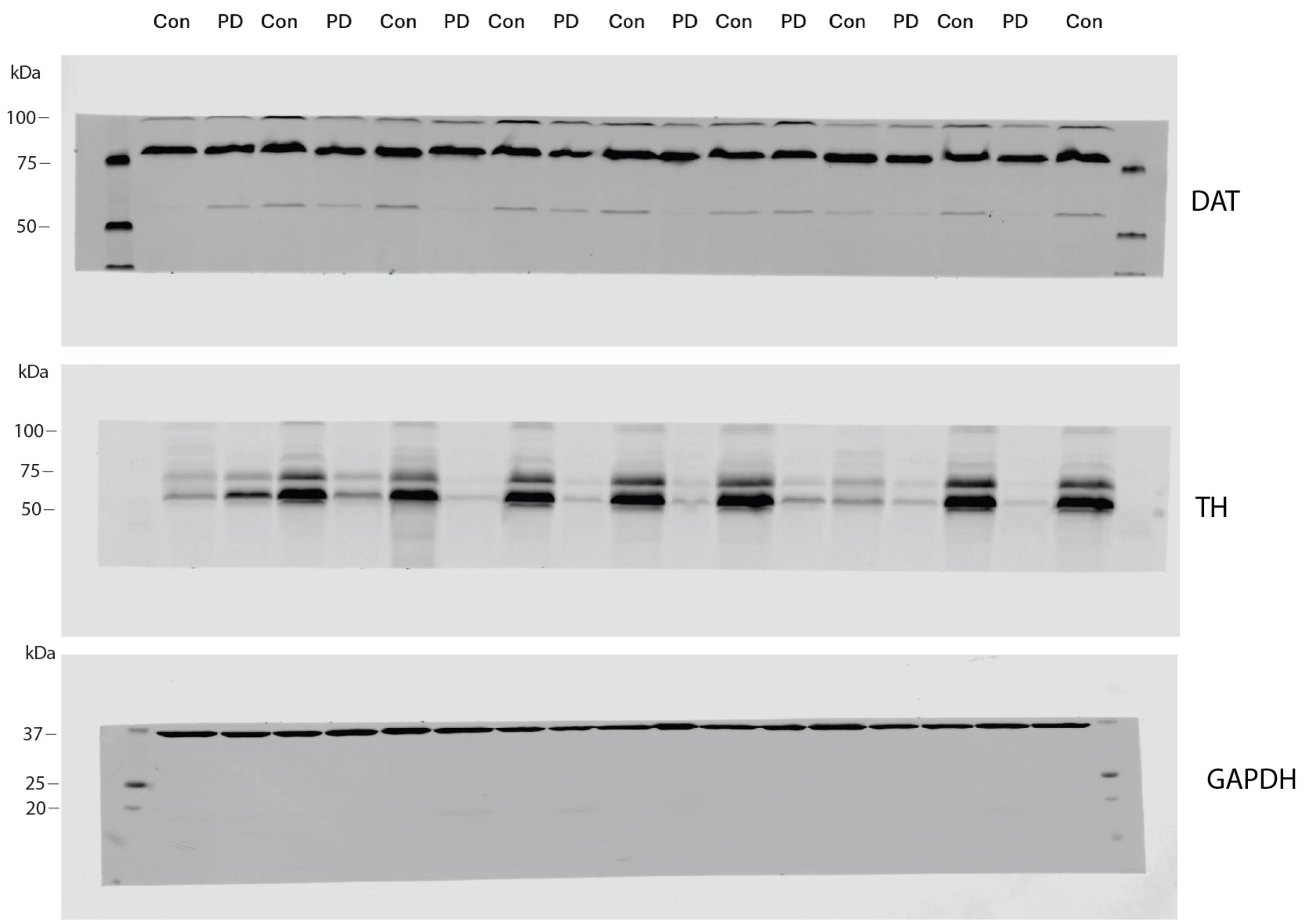


**Supplemental Fig. 9**: Uncropped Western blots corresponding to Figure 9 in main manuscript, showing dopamine transporter (DAT), tyrosine hydroxylase (TH), and GAPDH loading control in the substantia nigra of postmortem human tissue from neurologically intact controls (con) and Parkinson's disease (PD) brains.

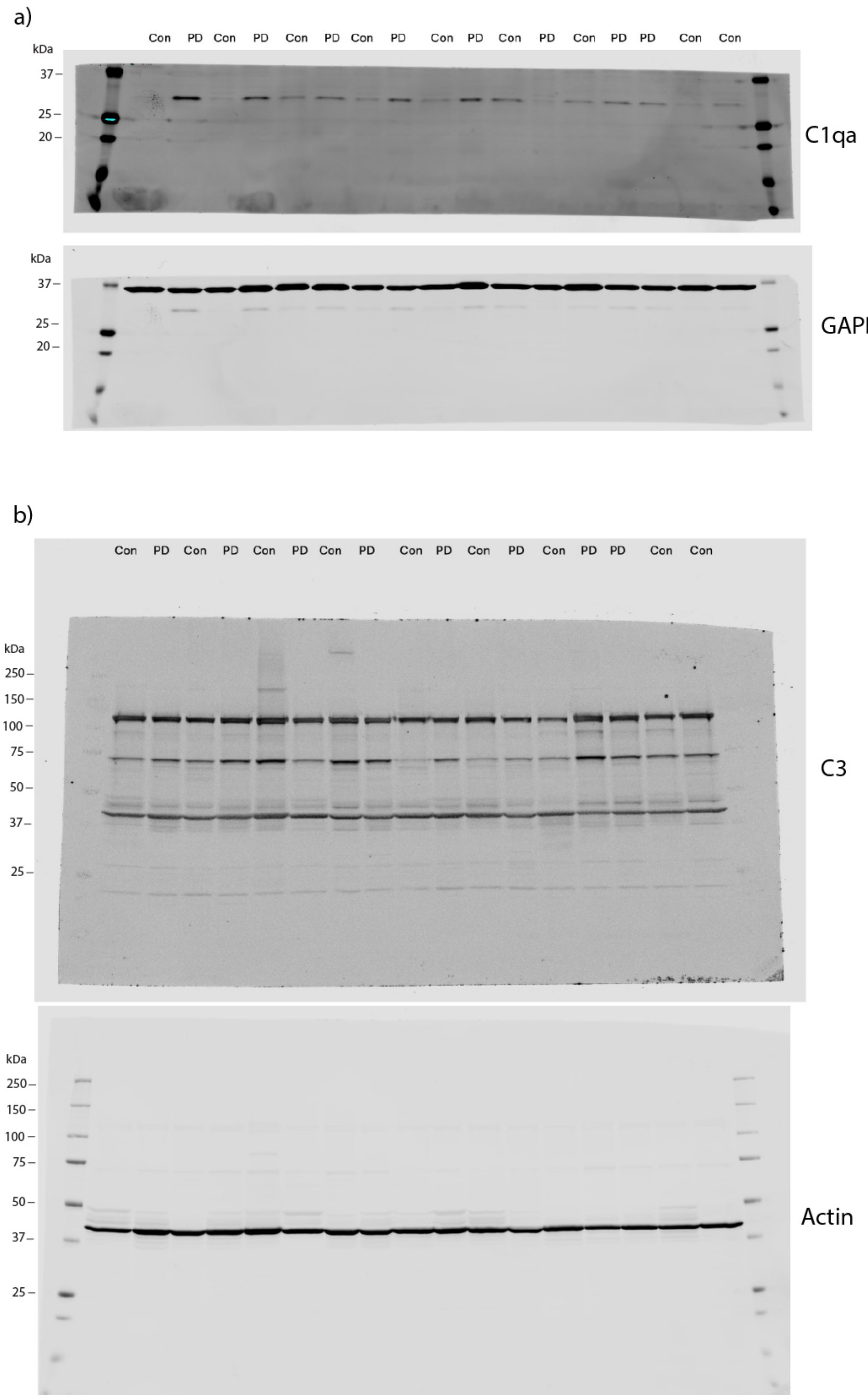


**Supplemental Fig. 10**: Uncropped Western blots corresponding to Figure 9 in main manuscript, showing C1qa (a), complement C3 (b), and associated loading controls in substantia nigra of postmortem human tissue from neurologically intact controls (con) and Parkinson’s disease (PD) brains.

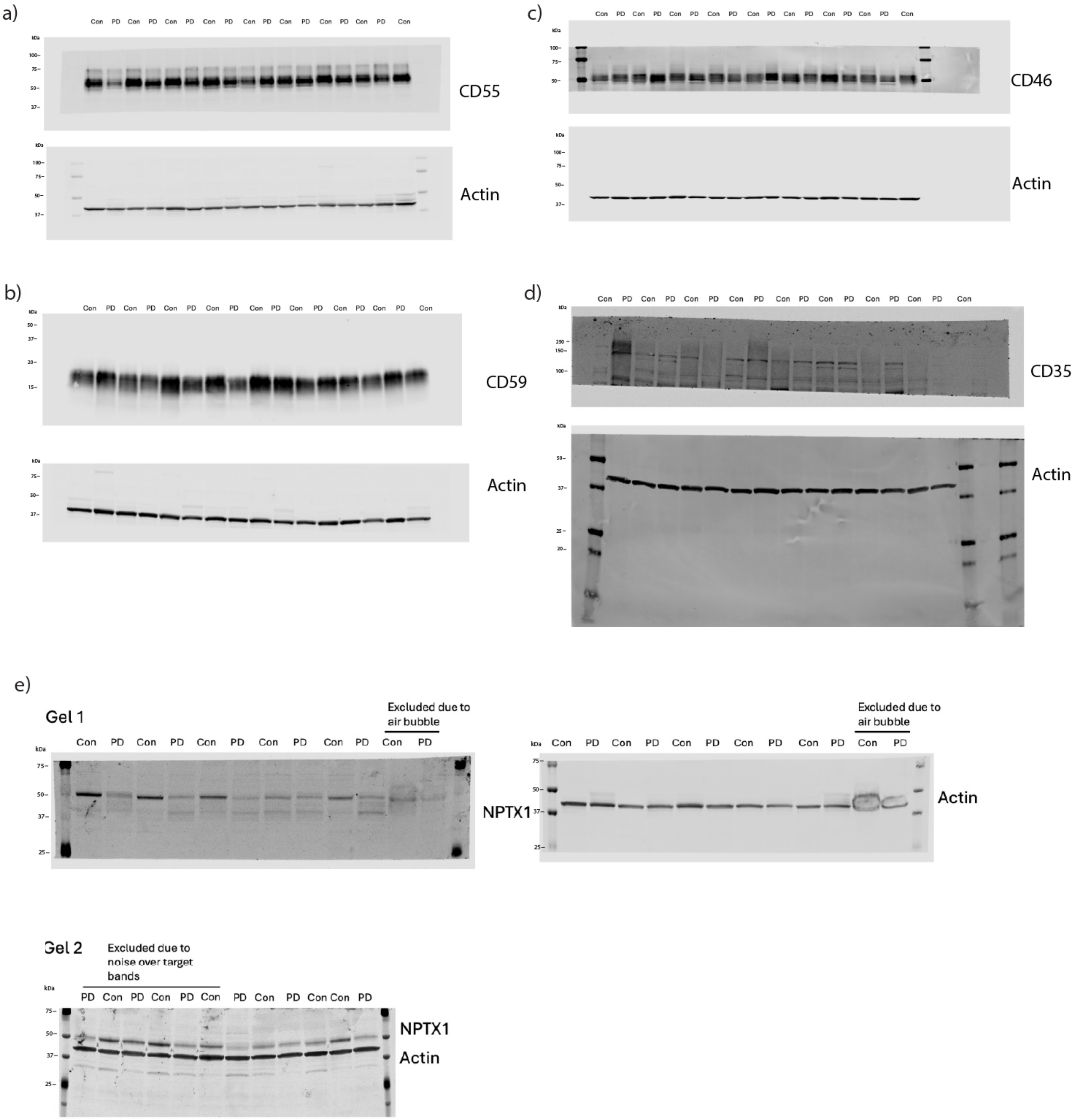


**Supplemental Fig. 10**: Uncropped Western blots corresponding to Figure 9 in main manuscript, showing CD55 (a), CD59 (b), CD46 (c), CD35 (d), and neuronal pentraxin 1 (NPTX1; e) and associated loading controls in substantia nigra of postmortem human tissue from neurologically intact controls (con) and Parkinson's disease (PD) brains.
